# Cooperative actin filament nucleation by the Arp2/3 complex and formins maintains the homeostatic cortical array in Arabidopsis epidermal cells

**DOI:** 10.1101/2022.08.03.502536

**Authors:** Liyuan Xu, Lingyan Cao, Jiejie Li, Christopher J. Staiger

## Abstract

Precise control over how and where actin filaments are created leads to the construction of unique cytoskeletal arrays within a common cytoplasm. Actin filament nucleators are key players in this activity and include the conserved Actin-Related Protein 2/3 (Arp2/3) complex, that creates dendritic networks of branched filaments, as well as a large family of formins that typically generate long, unbranched filaments and bundles. In some eukaryotic cells, these nucleators compete for a common pool of actin monomers and loss of one favors the activity of the other. To test whether this is a common mechanism, we combined the ability to image single filament dynamics in the homeostatic cortical actin array of living Arabidopsis (*Arabidopsis thaliana*) epidermal cells with genetic and/or small molecule inhibitor approaches to stably or acutely disrupt nucleator activity. We found that Arp2/3 mutants or acute CK-666 treatment markedly reduced the frequency of side-branched nucleation events as well as overall actin filament abundance. We also confirmed that plant formins contribute to side-branched filament nucleation in vivo. Surprisingly, simultaneous inhibition of both classes of nucleator increased overall actin filament abundance and enhanced the frequency of *de novo* nucleation events by an unknown mechanism. Collectively, our findings suggest that multiple actin nucleation mechanisms cooperate to generate and maintain the homeostatic cortical array of plant epidermal cells.

## Introduction

The actin cytoskeleton comprises a dynamic network of filaments that participates in a wide variety of cellular activities, including vesicle/organelle trafficking, cell expansion, cellulose synthesis, tissue/organelle development, and resistance to microbial pathogen invasion (Gao et al., 2009; Sampathkumar et al., 2013; Peremyslov et al., 2015; Scheuring et al., 2016; Qi and Greb, 2017; Li and Staiger, 2018; Zhang et al., 2019). Plant cells rearrange their actin cytoskeleton organization on timescales of seconds to minutes in response to biotic or abiotic stimuli, which means the actin cytoskeleton organization is extremely dynamic but also precisely controlled (Staiger et al., 2009; Henty-Ridilla et al., 2013b). Regulation of cytoskeletal organization and dynamics involves dozens of actin-binding proteins that participate in filament nucleation, elongation, severing, and depolymerization. Nucleation factors or nucleators initiate the formation of new actin filaments at a much faster rate than spontaneous nucleation from free actin monomers and are therefore considered key regulators of array formation and actin dynamics (Kadzik et al., 2020; Rosenbloom et al., 2021). Previous in-vitro studies, primarily from non-plant systems, have identified and characterized two classes of conserved actin nucleator, the Actin-Related Protein 2/3 (Arp2/3) complex and formins, with different specific activation mechanisms (Kovar, 2006; Pollard, 2007; Paul and Pollard, 2008; Burke et al., 2014; Suarez et al., 2015).

The Arp2/3 complex is a seven-subunit protein complex responsible for generating branched actin filaments and forming dendritic actin networks in eukaryotic cells (Mullins et al., 1998; Pollard, 2007). The ARP2 and ARP3 subunits mimic the structure of actin monomers. Once the Arp2/3 complex is activated, the ARP2 and ARP3 subunits along with a recruited actin monomer serve as the actin nucleus for the polymerization of a new (daughter) filament, which grows from the side of a pre-existing (mother) actin filament with a characteristic 70° angle (Mullins et al., 1998; Dayel and Mullins, 2004; Pollard, 2007; Beltzner and Pollard, 2008; Rouiller et al., 2008). This branched actin network structure is critical for many cellular processes in yeast and animal cells, such as cell division, cell migration, exocytosis, endocytosis and tissue or embryo development (Roh-Johnson and Goldstein, 2009; Cabrera et al., 2011; Sun et al., 2013; van der Kammen et al., 2017). In contrast, plants, such as Arabidopsis, with mutations in Arp2/3 subunits have relatively normal morphology and fertility; however, *arp2/3* mutants exhibit defects in epidermal cell morphology including distorted trichomes, cell-cell adhesion defects, and fewer lobes on leaf pavement cells (Szymanski et al., 1999; Le et al., 2003; Li et al., 2003; Mathur et al., 2003a; Harries et al., 2005; Yanagisawa et al., 2015). Recent research shows that the plant Arp2/3 complex is also involved in auxin transport (Peng et al., 2017; Sahi et al., 2018; García-González et al., 2020), guard cell opening and closing (Jiang et al., 2012; Li et al., 2014), resistance to penetration-mediated infection by fungal pathogens (Qin et al., 2021), and possibly in autophagy (Wang et al., 2016; Wang et al., 2019). Many of these studies reveal that *arp2/3* mutant cells have reduced actin filament abundance or misaligned actin bundles, suggesting that the functions of the Arp2/3 complex may be accomplished through generating specific actin structures. In growing trichome branches, for example, the Arp2/3 complex generates a tip-localized actin filament array constrained by cortical microtubules that coordinates growth (Yanagisawa et al., 2015; Yanagisawa et al., 2018). During fungal invasion, the Arp2/3 complex localizes to the site of penetration peg attack in epidermal pavement cells and generates a cortical actin filament patch necessary to suppress invasion (Qin et al., 2021). However, the molecular mechanisms by which the Arp2/3 complex coordinates the organization of cortical actin filament arrays in unstimulated cells or how it generates unique arrays in response to biotic and abiotic stress are poorly understood.

Formins represent another family of evolutionarily-conserved actin filament nucleators; they typically generate long linear actin filaments that can bundle together to form actin cables (Kovar, 2006; Pollard, 2007; Paul and Pollard, 2008). These actin cables provide tracks for intracellular organelle/vesicle trafficking. Formins have two universally conserved formin-homology (FH) domains, FH1 and FH2. The proline-rich FH1 domain allows formins to recruit profilin-bound actin to elongate actin filaments at the barbed ends. The FH2 domain, on the other hand, associates with other regions within itself for autoinhibition or it can interact with the FH2 domain of another formin to form a dimer that interacts with and suitably positions two actin monomers, thereby initiating the nucleation of linear actin filaments (Cvrčková et al., 2000; Deeks et al., 2002; Cheung and Wu, 2004; Cvrčková et al., 2004; Kovar, 2006; Michelot et al., 2006; Pollard, 2007; Paul and Pollard, 2008). There are 21 formin homologs in Arabidopsis and these are separated into two phylogenetic subclasses (Cvrčková et al., 2000). Based on sequence prediction, Class I formins have a signal peptide and an N-terminal transmembrane domain that target them to the plasma membrane, whereas Class II formins are predicted to have more diverse domain organization (Cvrčková et al., 2000; Deeks et al., 2002; Cheung and Wu, 2004; Cvrčková et al., 2004; Michelot et al., 2006). In addition, plant formin orthologs can be categorized according to the ability to associate with filament barbed ends and elongate growing filaments. Processive formins remain attached to the actin filament barbed end and move processively as the filament elongates, whereas non-processive formins remain at the filament nucleation site (Cvrčková et al., 2000; Deeks et al., 2002; Cheung and Wu, 2004; Cvrčková et al., 2004; Michelot et al., 2005; Michelot et al., 2006; Zhang et al., 2016). Biochemical results reveal that a non-processive Class I formin, FORMIN1 (AFH1), attaches to the side of a pre-existing actin filament to nucleate new filaments, suggesting a far more complicated role for formins in plant cells than just the ability to generate linear filaments (Michelot et al., 2006). In addition, both in-vivo and in-vitro experiments show that formins are critical for maintaining homeostatic actin array organization as well as regulating polarized cell growth (Vidali et al., 2009; Rosero et al., 2013; Lan et al., 2018). However, the detailed molecular mechanism of how formins regulate cytoskeletal organization and single actin filament dynamics in plant cells remains unclear.

The decision of when and where to generate an actin array as well as its specific structure and dynamics is essential for each actin array to specifically choreograph its cellular function and often requires coordination between the Arp2/3 complex and formins. Studies in fission yeast and some animal cells indicate that the Arp2/3 complex and formins compete for a limited supply of actin monomers and this competition prevents excessive activity of one or the other, thereby allowing cells to generate distinct actin structures and dynamics by regulating the balance of activities between these two nucleators (Hotulainen and Lappalainen, 2006; Lomakin et al., 2015; Suarez et al., 2015; Davidson et al., 2018; Antkowiak et al., 2019; Chan et al., 2019; Kadzik et al., 2020). During actin array assembly, the Arp2/3 complex generates branched filament arrays, whereas formins produce linear filament bundles (Carlier and Shekhar, 2017). Furthermore, in-vitro studies reveal that filaments generated by formins grow faster than Arp2/3-nucleated filaments (Vavylonis et al., 2006; Michelot et al., 2013; Suarez et al., 2015; Funk et al., 2019). Studies with M2 melanoma cells and HeLa cells illustrate that actin arrays generated by different nucleators have distinct properties of filament abundance, filament length, and turnover kinetics, but also reveal that both nucleators contribute to maintenance and turnover of the homeostatic actin cortex (Fritzsche et al., 2016). A recent study in Arabidopsis shows that actin arrays in cotyledon epidermal pavement cells are slightly more dense and the extent of bundling is unchanged when either a Class I formin, AFH1, or the Arp2/3 subunit, ARPC5, are genetically down-regulated; however, single filament dynamics or nucleation events were not evaluated directly (Cifrová et al., 2020). The authors propose that the Arp2/3 complex and formins in plants may have complementary roles to overcome the loss of the other. Therefore, it is important to understand how each nucleator regulates the structure and dynamic properties of actin arrays individually to better understand how they contribute to the cortical actin homeostasis and how they are coordinated to regulate the actin cytoskeleton during different biological processes.

Quantitative analysis of single actin filament dynamics in vivo is relatively hard to conduct in yeast and mammalian cells, because either the cells are too small or the actin arrays are too dense and filament lengths are below the limits of resolution of light microscopy. Epidermal cells from dark-grown Arabidopsis hypocotyls are large in size and have relatively sparse actin arrays in the cortical cytoplasm, making this a powerful model system for quantitative analysis of single actin filament dynamics, especially when combined with high spatial and temporal resolution imaging approaches and genetically-encoded fluorescence reporters (Staiger et al., 2009; Henty et al., 2011; Henty-Ridilla et al., 2013a; Cai et al., 2014; Cao et al., 2016; Arieti and Staiger, 2020). Previous work demonstrates a remarkably high rate of new filament construction, rapid growth at filament ends, and disassembly by prolific severing activity in a mechanism termed “stochastic dynamics” (Staiger et al., 2009). High rates of polymerization are likely supported by a large pool of monomeric actin that is buffered with an excess of the monomer-binding protein, profilin, to suppress spontaneous nucleation events (Chaudhry et al., 2007; Staiger et al., 2009; Cao et al., 2016). The ability to observe both actin filament architecture and activities of single filaments in living cells allows direct visualization and quantitative analysis of the effects associated with the disruption of actin nucleators on filament activities in vivo (Cao et al., 2016).

In this study, we test the hypothesis that the Arabidopsis Arp2/3 complex mediates nucleation of side-branched filaments in vivo but does not facilitate filament elongation directly. We used both genetic and pharmacological approaches to disrupt Arp2/3 complex activity in plant cells. With quantitative live-cell imaging at single filament resolution, we demonstrated directly that the Arp2/3 complex is responsible for the nucleation of side-branched filaments in vivo. We also characterized the difference between the Arp2/3 complex and formins with respect to the dynamic behaviors of the filaments they generate and evaluated the consequence of losing both classes of filament nucleator on actin organization and dynamics. Surprisingly, living plant cells deficient for two classes of nucleator generate comparatively normal and dynamic actin arrays by an apparent *de novo* filament nucleation mechanism.

## Results

### Arabidopsis plants have defects in general growth and epidermal cell morphology when deficient for the Arp2/3 complex

Previous studies of Arp2/3 complex mutants reveal defects in Arabidopsis epidermal cell growth and morphology (Szymanski et al., 1999; Le et al., 2003; Li et al., 2003; Mathur et al., 2003b; Harries et al., 2005; Facette et al., 2015; Yanagisawa et al., 2015; Cifrová et al., 2020). Here, we acquired a T-DNA insertion mutant for *ARP2*, *arp2-1* (or *wrm1-2*; SALK_003448), and a point mutation of *ARPC2*, *arpc2* (or *dis2-1*; El-Assal et al., 2004), and confirmed the cell and organ growth phenotypes. We observed that both *arp2-1* and *arpc2* homozygous mutant plants had adhesion defects at end walls of hypocotyl epidermal cells and severely distorted leaf trichomes when compared to wild-type sibling lines (Supplemental Fig. S1A). In addition, etiolated hypocotyls from *arp2-1* and *arpc2* seedlings were significantly shorter than wild-type seedlings at the same time points (Supplemental Fig. S1, B and C). To test whether these growth defects resulted from defects in cell expansion, we examined epidermal cell length and found that cells in all regions of etiolated *arp2/3* hypocotyls were significantly shorter compared to wild-type hypocotyls; however, cell widths were not affected (Supplemental Fig. S1, D and E). Similar results for root growth were observed in light-grown seedlings (Supplemental Fig. S1, F and G). These results confirm that the Arp2/3 complex plays a role in axial cell expansion in both dark-grown hypocotyls and light-grown roots of Arabidopsis.

### Actin filament abundance and bundling are reduced in *arp2/3* mutants

To assess the influence of the Arp2/3 complex on cortical actin cytoskeletal organization, we compared images of epidermal cells from the apical region of 5-d-old etiolated wild-type, *arp2-1,* and *arpc2* hypocotyls collected by variable-angle epifluorescence microscopy (VAEM) (Staiger et al., 2009). Homozygous mutant and isogenic wild-type sibling lines that express the actin reporter, green fluorescent protein (GFP) fused with FIMBRIN1 actin-binding domain 2 or GFP-fABD2 (Sheahan et al., 2004) were prepared by crossing to facilitate the observation and measurement of actin cytoskeleton organization and filament dynamics (Staiger et al., 2009). The actin array structure in *arp2-1* and *arpc2* cells had fewer, thinner, and more scattered filaments compared to corresponding wild-type cells (Fig. 1A). To compare these differences quantitively, we used *density*, *skewness* and *the coefficient of variation (CV)* as three key parameters established previously for standardized descriptions of actin cytoskeleton array organization in living cells (Higaki et al., 2010; Ueda et al., 2010; Henty et al., 2011; Higaki et al., 2020). Density measures the percentage of occupancy of actin filaments in an array, whereas both skewness and CV define the extent of bundling of actin filaments. CV uses a different calculation method compared to skewness, and is reportedly a better indicator of bundling, especially for VAEM images (Higaki et al., 2020). Actin density in *arp2-1* and *arpc2* cells was significantly decreased compared to the respective wild-type cells (Fig. 1B), and the extent of filament bundling was significantly reduced as well when measured by skewness or CV methods (Fig. 1C and D). These results demonstrate that the Arp2/3 complex contributes to the generation of both individual filaments and filament bundles and is necessary for maintaining the homeostatic organization of the cortical actin cytoskeleton in epidermal cells.

**Figure 1.**
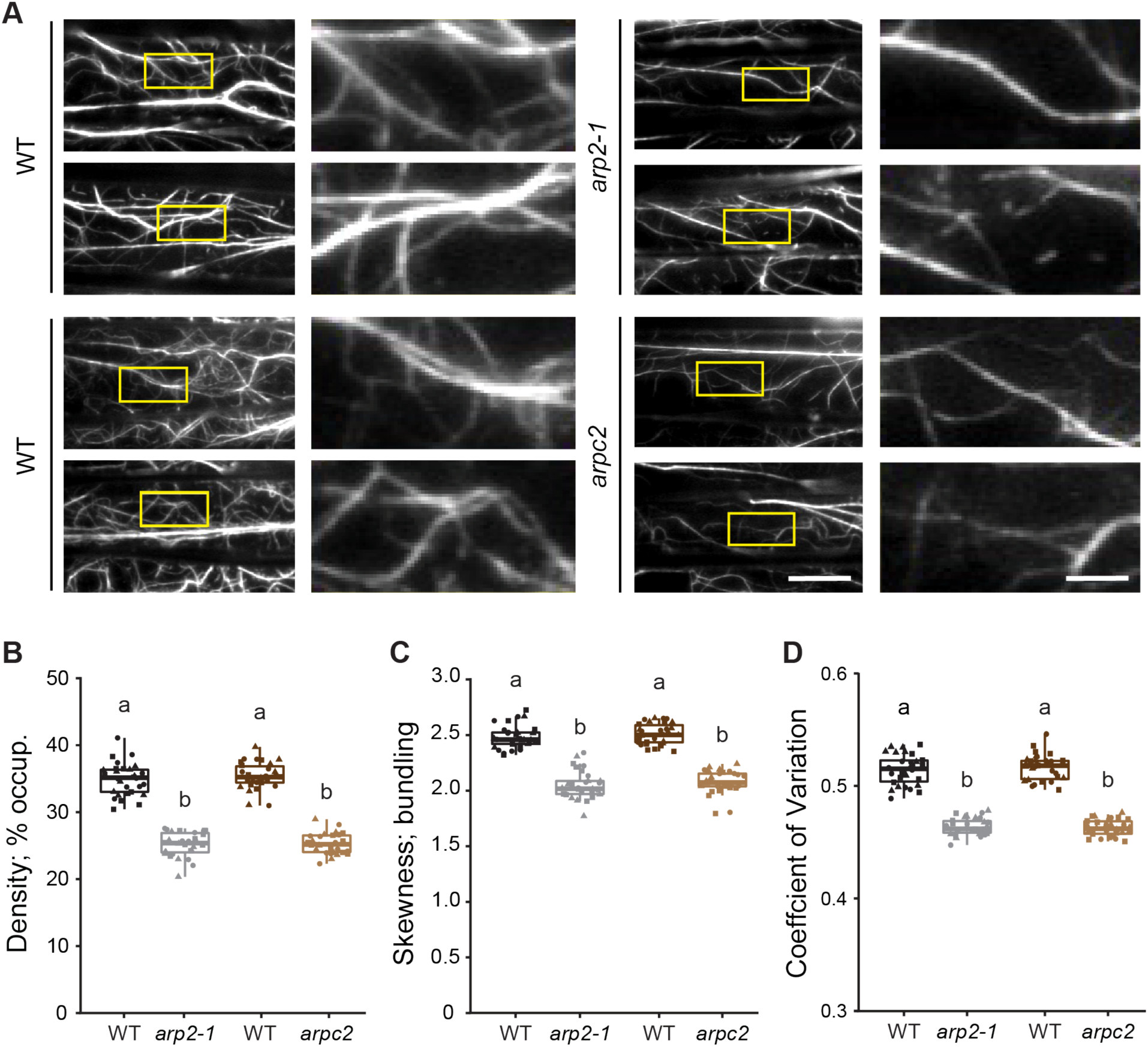
Genetic disruption of the Arp2/3 complex leads to reduced actin filament density and bundling. **A)** Representative images of epidermal cells from the apical region of 5-d-old etiolated hypocotyls expressing GFP-fABD2 imaged by variable angle epifluorescence microscopy (VAEM) are shown in left columns. Scale bar: 20 μm. Regions of interest (yellow boxes) were magnified and shown in right columns. Scale bar: 5 μm. **B–D)** Quantitative analysis of the percentage of occupancy or density of actin filament arrays **(B)** and the extent of filament bundling as measured by skewness **(C)** and coefficient of variance **(D)** analyses. Both the density and the bundling of actin arrays in *arp2-1* and *arpc2* cells were significantly decreased compared to those in the respective wild-type cells. In box-and-whisker plots, boxes show the interquartile range (IQR) and the median, and whiskers show the maximum-minimum interval of three biological repeats with independent populations of plants. Individual biological repeats are represented with different shapes (n = 30 seedlings, 10 seedlings per biological repeat). Letters [a–b] denote groups that show statistically significant differences with other genotypes by one-way ANOVA with Tukey’s post-hoc test (P < 0.05).

### The Arp2/3 complex plays a significant role in filament nucleation and generation of side-branched filaments

To investigate whether the Arp2/3 complex participates in actin filament nucleation, we measured single actin filament formation and dynamics in wild-type and *arp2/3* mutant cells by collecting time-lapse movies from the apical regions of 5-d-old dark-grown hypocotyls with VAEM. The sparse nature of the cortical actin array in hypocotyl epidermal cells and the high spatial and temporal resolution afforded by VAEM allowed us to visualize (Fig. 2A) and quantify individual nucleation events with a previously described assay (Cao et al., 2016). Briefly, we counted all new or regrowing filaments identified in multiple 400 µm^2^ regions of interest (ROI) within a cell during a 100-s time-lapse movie. The nucleation frequency was normalized to the average filament number within each ROI to minimize the influence of filament abundance differences between ROIs and genotypes. The overall nucleation frequency in *arp2-1* cells (0.38 ± 0.03 events/filament/min) was significantly decreased compared to wild-type cells (0.62 ± 0.05 events/filament/min) (Fig. 2B; Supplemental Movie S1; Supplemental Movie S2) and coincided with our observation that loss of function of the Arp2/3 complex resulted in a significant reduction in filament abundance (Fig. 1B). Similar reductions in overall nucleation frequency were also observed for *arpc2* compared to wild-type cells (Supplemental Fig. S2A). Next, we classified nucleation events into three different subpopulations based on filament origin: *de novo* from the cytoplasm; from the side of a pre-existing filament or bundle; or, from the end of a pre-existing filament (Fig. 2A; Supplemental Movies S3 to S5) (Staiger et al., 2009; Henty-Ridilla et al., 2013a; Cao et al., 2016). Only the side-branched nucleation frequency was significantly decreased in *arp2-1* and *arpc2* cells compared to the respective wild-type sibling cells (Fig. 2C– E; Supplemental Movies S1 and S2; Supplemental Fig. S2, C–E). Side-branched nucleation events contribute about half of all new filament origins and these were reduced by 60–70% in the *arp2/3* mutants.

**Figure 2.**
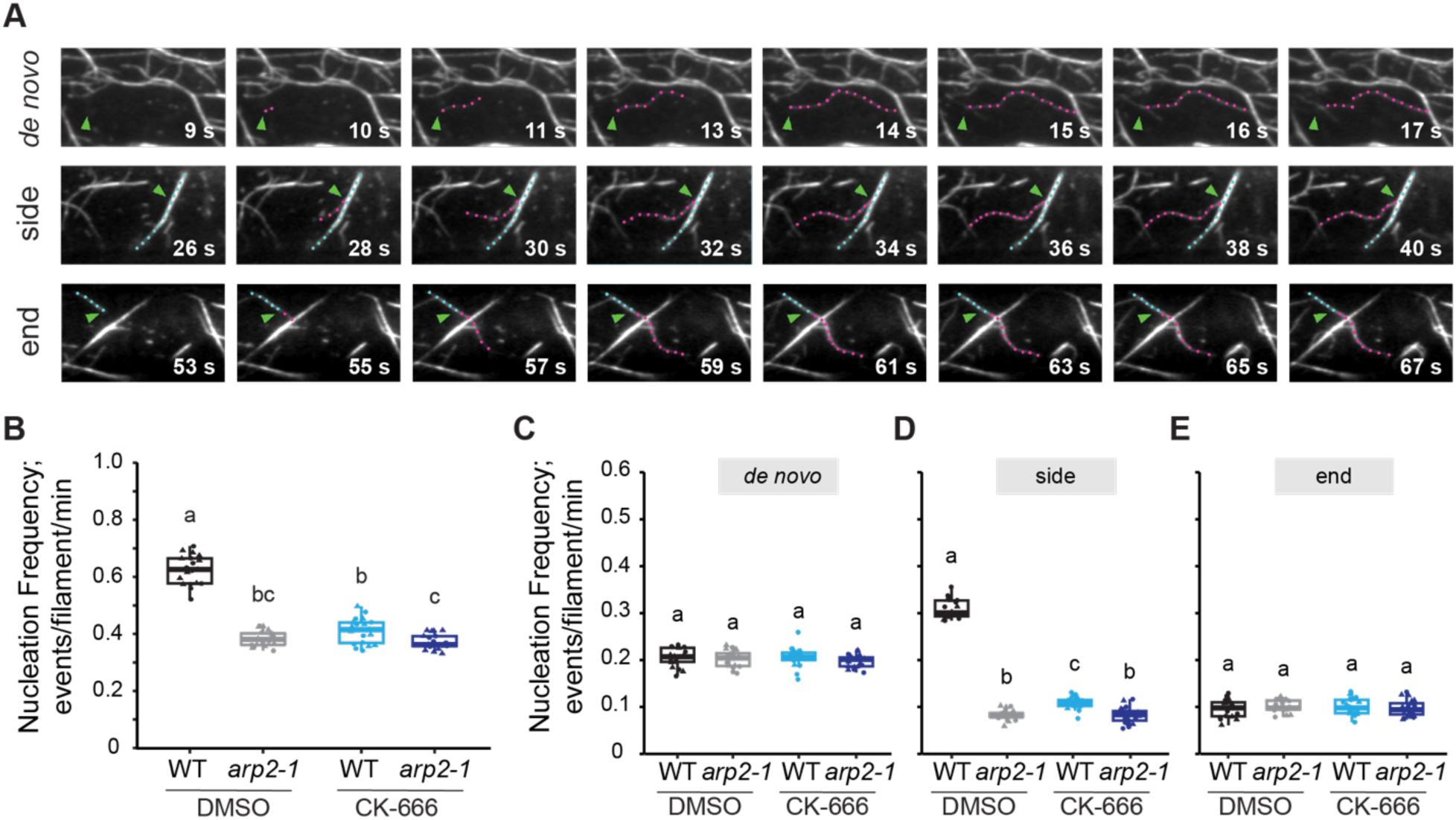
Actin filament nucleation frequency is decreased by chemical or genetic inhibition of the Arp2/3 complex. **A)** Representative time-lapse series showing three subclasses of actin filament origin identified in hypocotyl epidermal cells: Actin filaments initiated *de novo* in the cytoplasm (top), from the side of a pre-existing filament (middle), or from the end of a pre-existing filament (bottom row). Blue dots; pre-existing filament. Magenta dots; new growing filament. Green arrowheads; nucleation site. Scale bar: 5 μm. **B–E)** Quantitative analysis of actin filament nucleation frequency, both overall **(B)** and by subclass of origin **(C–E)**. Hypocotyls were treated with DMSO solution (0.05% DMSO) or 10 µM CK-666 for 5 min prior to imaging with VAEM. The overall nucleation frequency **(B)** for each genotype or treatment was defined as the total number of filament origins per filament per minute in a 400 μm^2^ region of interest. Total nucleation frequency in DMSO-treated wild-type cells was significantly higher than DMSO-treated *arp2-1*, CK-666-treated wild-type, or CK-666-treated *arp2-1* cells. When filament origin events were categorized into *de novo* **(C)**, side **(D)**, and end populations **(E)**, only the side-branching nucleation events showed a significant reduction in *arp2-1* and CK-666-treated cells compared to DMSO-treated wild type. In box-and-whisker plots, boxes show the interquartile range (IQR) and the median, and whiskers show the maximum-minimum interval of two biological repeats with independent populations of plants. Individual biological repeats are represented with different shapes (n = 20 seedlings, 10 seedlings per biological repeat). Letters [a–c] denote groups that show statistically significant differences with other genotypes or treatments by two-way ANOVA with Tukey’s post-hoc test (P < 0.05).

A previous study reports different results regarding the actin organization in *arp2/3* mutants; instead of significantly decreased filament abundance as observed here, a minor increase in actin filament density in *arpc5* cotyledon pavement cells was reported (Cifrová et al., 2020). These differences could result from the use of a different *arp2/3* complex subunit mutation, the investigation of epidermal cells from a different tissue, the choice of actin cytoskeleton reporter, or all of the above. Several previous publications show that LifeAct reduces both the actin reorganization rate and actin polymerization rate (van der Honing et al., 2011; Spracklen et al., 2014; Courtemanche et al., 2016), and displays longer and thicker filaments (Flores et al., 2019). To test which factors led to differing results between the current work and that of Cifrová et al. (2020), we conducted a direct comparison between the ability of GFP-fABD2 or GFP-LifeAct to report actin architecture and single filament dynamics in either etiolated hypocotyls (Supplemental Fig. S3) or light-grown cotyledons (Supplemental Fig. S4). Moreover, we introduced GFP-LifeAct into *arp2-1* to compare directly with the GFP-fABD2 *arp2-1* reporter line described above. Our results showed that *arp2-1* expressing either GFP-fABD2 or GFP-LifeAct had significantly lower actin filament density in the cortical array of both etiolated hypocotyl and light-grown cotyledon epidermal cells (Supplemental Fig. S3B and S4B). Similarly, overall actin filament nucleation frequency and the side-branched nucleation subclass were significantly reduced in *arp2-1* epidermal cells from both reporter lines, in either hypocotyls or cotyledons, compared to the corresponding wild-type lines (Supplemental Fig. S3, E–H; Supplemental Fig. S4, E–H). These results suggest that using GFP-fABD2 as the actin reporter, or imaging epidermal cells of the etiolated hypocotyl, are not causal for the actin phenotypes we observed in *arp2/3* mutants. Because it is beyond the scope of the current investigation, we did not seek to determine whether the previous findings are specific to the *arpc5* mutant allele used, but note that we find similar quantitative differences in actin architecture and nucleation frequency with mutations in two different subunits (*arp2-1* and *arpc2*) of the Arp2/3 complex. Considering the nature of the LifeAct marker in stabilizing actin structures and our primary goal of observing and analyzing the dynamic behaviors of single actin filaments, we used the GFP-fABD2 marker lines for the remainder of the experiments in this study.

### Actin filaments generated by the Arp2/3 complex have unique dynamic properties

To understand how the Arp2/3 complex influences the dynamics of actin filaments, we tracked many dozens of individual filaments from their first appearance to complete disappearance and measured several parameters that were previously established to describe actin filament turnover (Fig. 3A; Table 1; Supplemental Movies S6 and S7)(Staiger et al., 2009; Henty et al., 2011). Compared to wild type, the population distribution of actin filament elongation rates in *arp2-1* cells was more left skewed, and the average filament elongation rate was also higher (Fig. 3B; Table 1), indicating that actin filaments were growing significantly faster in *arp2-1* cells. It is also obvious that there were two or three distinct populations of filament elongation rates with peaks at 0.75 – 1.25, 2.0 – 2.25, and > 3 µm/s with the latter two categories becoming more prevalent in *arp2-1* cells (Fig. 3B). Similar differences in overall elongation rate and fast-growing filament populations were observed in *arpc2* (Supplemental Table S1; Supplemental Fig. S2B). Filaments in both *arp2-1* and *arpc2* mutant cells also had significantly longer filament lengths and lifetimes than the ones in respective wild-type cells (Table 1; Supplemental Table S1).

**Figure 3.**
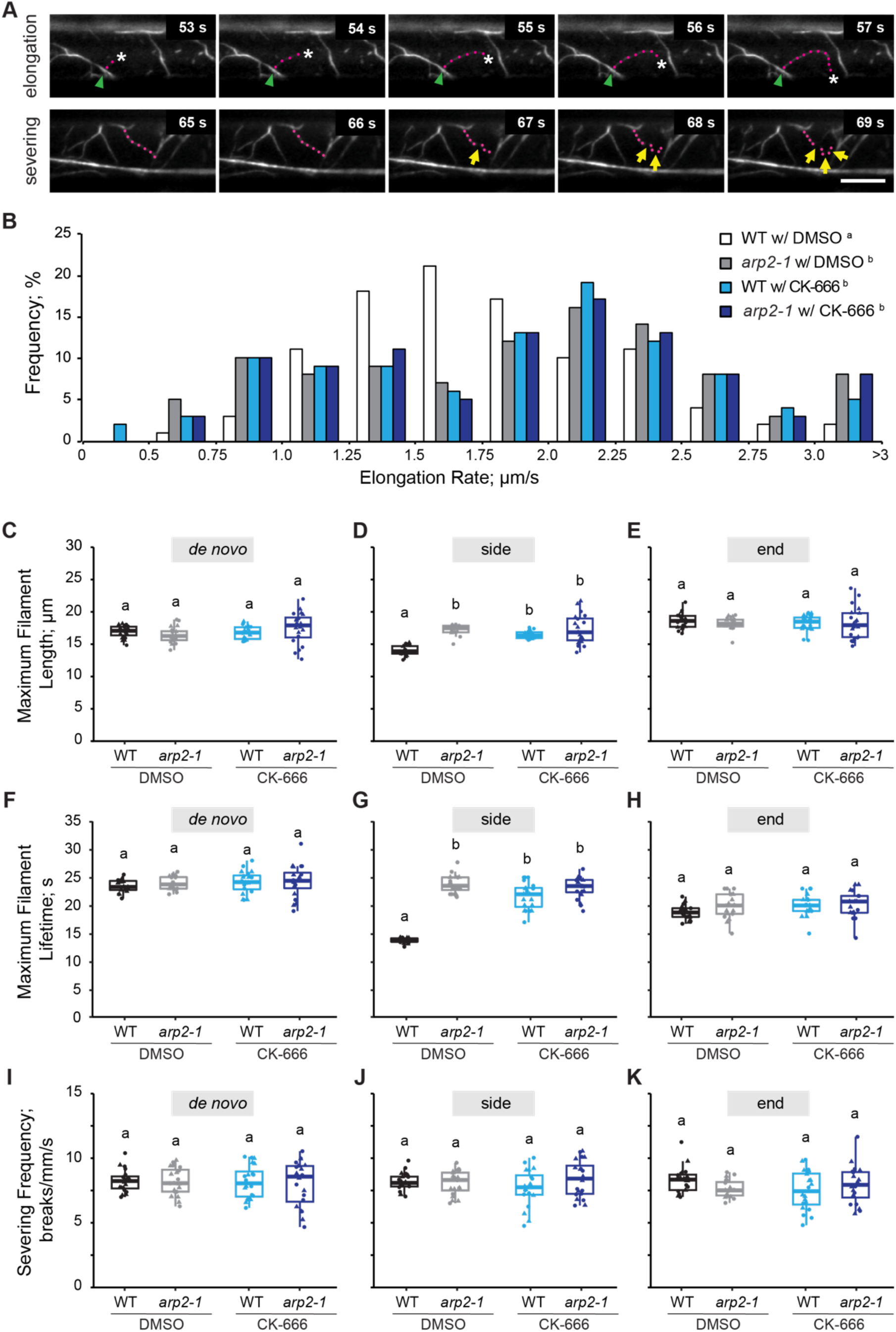
Actin filament dynamic properties are altered when the Arp2/3 complex is inhibited. **A)** A representative actin filament was tracked during the elongation phase (top) and another filament was fragmented into pieces during the severing phase (bottom). Magenta dots; growing actin filament. Green arrowhead; nucleation site. White asterisk; elongating filament end. Yellow arrows; severing events. Scale bar: 10 μm. **B)** Quantitative analysis of the population distribution and average elongation rate of actin filaments in hypocotyl epidermal cells. Hypocotyls were treated with 0.05% DMSO solution or 10 µM CK-666 for 5 min prior to imaging with VAEM. The elongation rate distribution of DMSO-treated wild type had a single peak at 1.25 – 1.75 μm/s, whereas the Arp2/3-inhibited groups had three peaks at 0.75 – 1.25 μm/s, 2.0 – 2.5 μm/s, and > 3 μm/s. n ≥ 100 single filaments from two individual biological repeats (for one biological repeat, 5 single filaments were counted in one hypocotyl from at least 10 hypocotyls per genotype or treatment). Letters [a–b] denote genotypes or treatments that show statistically significant differences with other groups by Chi-squared test, P < 0.05. **C–E)** The average maximum length of side-branching filaments in DMSO-treated wild-type cells was significantly shorter than that in DMSO-treated *arp2-1*, CK-666-treated wild-type, or CK-666-treated *arp2-1* cells; however, filaments that originated *de novo* or from pre-existing ends did not show any significant difference. **F–H)** The average maximum lifetime of side-branching filaments in DMSO-treated wild-type cells was significantly shorter than that in DMSO-treated *arp2-1*, CK- 666-treated wild-type, or CK-666-treated *arp2-1* cells; however, filaments that originated *de novo* or from pre-existing ends did not show any significant difference. **I–K)**, The severing frequency did not show any significant difference between different genotypes or treatments. For box-and-whisker plots, boxes show the interquartile range (IQR) and the median, and whiskers show the maximum-minimum interval of two biological repeats with independent populations of plants. Individual biological repeats are represented with different shapes (n = 20 seedlings, 10 seedlings per biological repeat). Letters [a–b] denote groups that show statistically significant differences with other genotypes or treatments by two-way ANOVA with Tukey’s post-hoc test (P < 0.05).

**Table 1.**
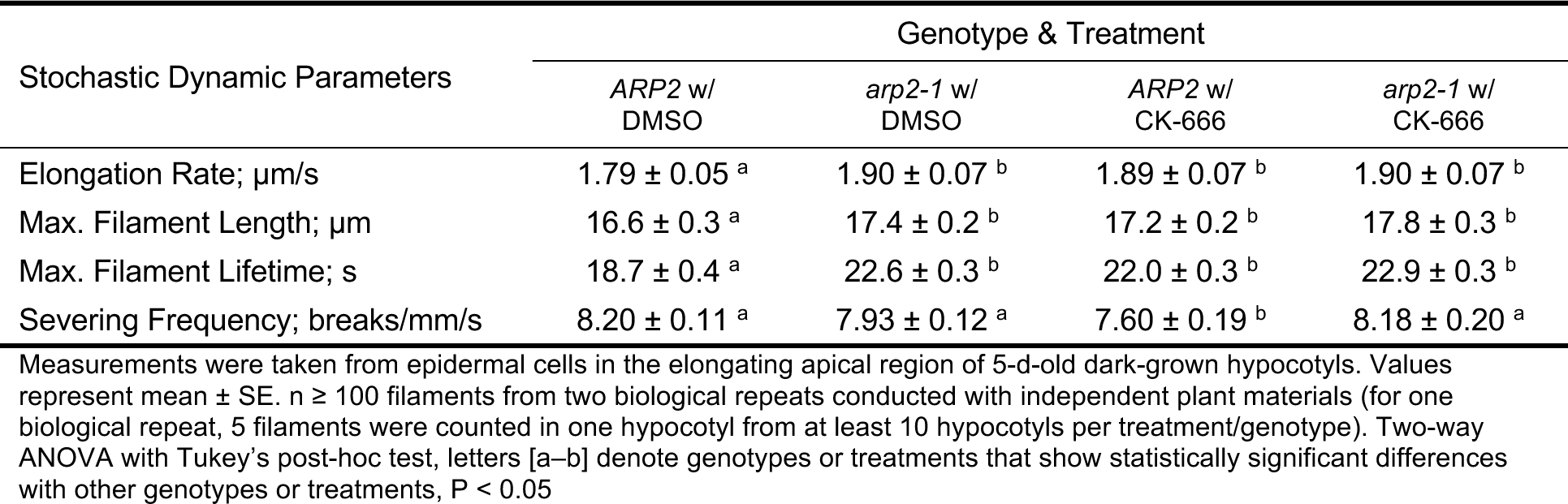
Single actin filament dynamics in wild type (*ARP2*) and *arp2-1* mutant with or without CK-666.

| Stochastic Dynamic Parameters | Genotype & Treatment |  |  |  |
| --- | --- | --- | --- | --- |
|  | *ARP2* w/<br>DMSO | *arp2-1* w/<br>DMSO | *ARP2* w/<br>CK-666 | *arp2-1* w/<br>CK-666 |
| Elongation Rate; $\mu\text{m/s}$ | $1.79 \pm 0.05^a$ | $1.90 \pm 0.07^b$ | $1.89 \pm 0.07^b$ | $1.90 \pm 0.07^b$ |
| Max. Filament Length; $\mu\text{m}$ | $16.6 \pm 0.3^a$ | $17.4 \pm 0.2^b$ | $17.2 \pm 0.2^b$ | $17.8 \pm 0.3^b$ |
| Max. Filament Lifetime; s | $18.7 \pm 0.4^a$ | $22.6 \pm 0.3^b$ | $22.0 \pm 0.3^b$ | $22.9 \pm 0.3^b$ |
| Severing Frequency; breaks/mm/s | $8.20 \pm 0.11^a$ | $7.93 \pm 0.12^a$ | $7.60 \pm 0.19^b$ | $8.18 \pm 0.20^a$ |

Measurements were taken from epidermal cells in the elongating apical region of 5-d-old dark-grown hypocotyls. Values represent mean ± SE. $n \geq 100$ filaments from two biological repeats conducted with independent plant materials (for one biological repeat, 5 filaments were counted in one hypocotyl from at least 10 hypocotyls per treatment/genotype). Two-way ANOVA with Tukey's post-hoc test, letters [a–b] denote genotypes or treatments that show statistically significant differences with other genotypes or treatments, $P < 0.05$

Previous in vitro data suggest that filament ends generated by the Arp2/3 complex and formins have different elongation rates; many formins are processive polymerases that use profilin-actin to add monomers to growing filaments (Kovar et al., 2003; Romero et al., 2004; Akin and Mullins, 2008; Zhang et al., 2016). To test whether dynamic properties of filaments with different nucleation patterns are distinct from each other, we measured the full suite of dynamic parameters for each population. We found that side-branched filaments in *arp2-1* cells were significantly longer (Fig. 3, C–E) and filament lifetime was prolonged (Fig. 3, F–H) compared to the side-branched filaments in wild-type cells; however, the severing frequency did not show a significant difference between wild-type and *arp2-1* cells (Fig. 3, I–K). These dynamic differences between side-branched filaments in *arpc2* mutant and wild-type sibling cells were conserved (Supplemental Fig. S2, C–K).

Collectively, these data demonstrate that the Arp2/3 complex is responsible for generating new filaments from the side of pre-existing filaments and is necessary for cells to maintain the homeostatic actin cytoskeleton architecture. Moreover, Arp2/3-nucleated filaments have distinct dynamic properties compared with those generated *de novo* or from the end of a pre-existing filament; in particular, they grow slower and are shorter, and have reduced filament lifetimes.

### The small molecule inhibitor CK-666 phenocopies actin-based defects in *arp2* and *arpc2* mutants

Several studies report that CK-666, a small molecule inhibitor of the Arp2/3 complex that is effective on yeast and animal cells (Nolen et al., 2009; Hetrick et al., 2013), phenocopies the effects of *arp2/3* mutants on tomato pollen tube growth (Liu et al., 2020) and *Arabidopsis* sperm nuclear migration (Ali et al., 2020), both of which depend upon the actin cytoskeleton. Moreover, a recent study demonstrates that CK-666 treatment does not influence actin redistribution between the basal and apical cell region in hypocotyl epidermal cells that occurs during early embryo growth (Cui et al., 2023). Because the earlier studies (Liu et al., 2020; Ali et al., 2020) did not directly demonstrate the effects of CK-666 on actin organization or filament nucleation, it is necessary to validate whether CK-666 influences the function of the Arp2/3 complex in plant cells, as a side-branched actin filament nucleator, before we use it as an effective plant Arp2/3 inhibitor. We hypothesized that applying CK-666 to wild-type Arabidopsis cells would inhibit the Arp2/3 complex activity and mimic the *arp2/3* mutant phenotypes described above. To test the effects of CK-666, we applied a dose series (0, 1, 5, 10, 50, and 100 µM) for 5 min and collected snapshots of epidermal cells from the apical region of treated hypocotyls (Supplemental Fig. S5). Compared to the mock treatment (0 µM), CK-666 showed a significant reduction in both filament density and the extent of bundling starting at 5 µM and reached its maximum effect at 50–100 µM (Supplemental Fig. S5, B–D). A time-course (0, 5, 10, 30, and 60 min) of 10 µM CK- 666 treatment showed that 5 min was sufficient to alter actin array organization (Supplemental Fig. S5, E–G). Finally, cells were able to recover normal actin array architecture from a 5 min, 10 µM CK-666 application after a 30-min wash out (Supplemental Fig. S5, H–J).

To further examine whether CK-666 phenocopies *arp2/3* mutants, we treated 5-d-old etiolated hypocotyls of wild type and *arp2-1* with either DMSO (mock) solution or 10 μM CK-666 for 5 min and measured actin organization and single filament dynamics. Compared to DMSO-treated wild-type cells, the CK-666-treated wild-type cells had significantly decreased filament abundance (Fig. 4B), extent of filament bundling (Fig. 4, C and D), as well as overall nucleation frequency (Fig. 2B), and these changes mirrored actin organization in DMSO-treated *arp2-1* cells. Moreover, treatment of *arp2-1* with CK-666 showed no differences in terms of filament organization or dynamics compared to *arp2-1* alone (Fig. 3 and 4). CK-666 treatment also potently suppressed side-branched filament nucleation in wild-type cells, but not end or *de novo* filament origins (Fig. 2, C–E; Supplemental Movies S1 and S8), thereby fully phenocopying the effects of *arp2-1*. Finally, treatment of *arp2-1* with CK-666 did not further reduce filament side-branch nucleation events, indicating that its likely mode of action is to inhibit daughter filament formation from a mother filament by the Arp2/3 complex (Fig. 2D; Supplemental Movies S8 and S9).

**Figure 4.**
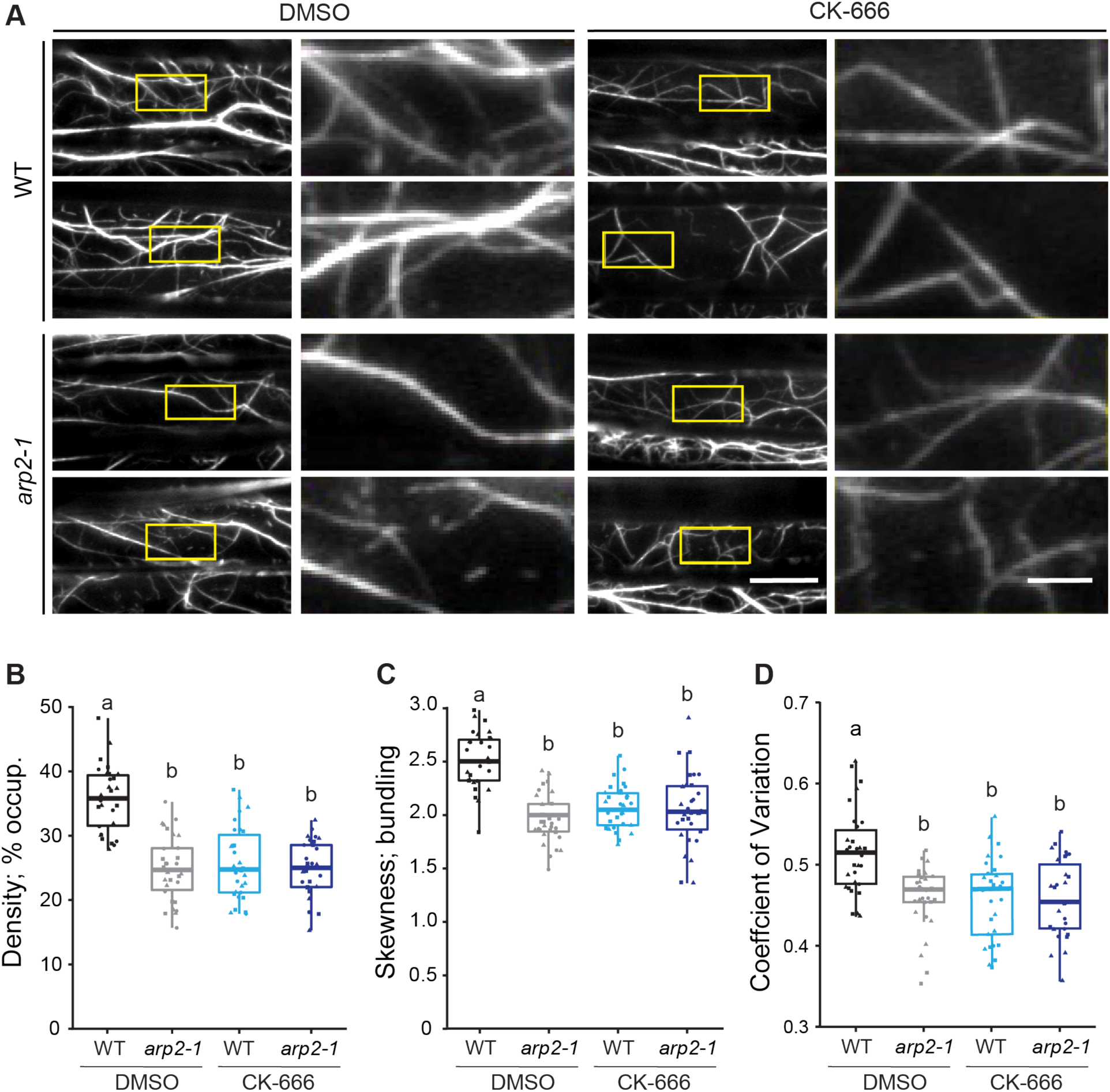
A small molecule inhibitor of the Arp2/3 complex, CK-666, reduces actin filament density and bundling. **A)** Representative images of epidermal cells from the apical region of 5-d-old etiolated hypocotyls are shown in left columns. Scale bar: 20 μm. Regions of interest (yellow boxes) were magnified and displayed in right columns. Scale bar: 5 μm. Hypocotyls were treated with 0.05% DMSO solution or 10 µM CK-666 for 5 min prior to imaging with VAEM. Actin filament arrays in DMSO-treated *arp2-1*, CK-666-treated *ARP2*, and CK-666-treated *arp2-1* cells appeared to be less dense and less bundled compared to DMSO-treated wild-type cells. **B–D)** Quantitative analysis of the percentage of occupancy or density of actin filament arrays **(B)**, and the extent of filament bundling as measured by skewness **(C)** and coefficient of variation **(D)** analyses. Both the density and bundling of actin arrays in DMSO-treated *arp2-1*, CK-666-treated wild-type and CK-666-treated *arp2-1* cells were significantly decreased compared to DMSO-treated wild-type cells. In box-and-whisker plots, boxes show the interquartile range (IQR) and the median, and whiskers show the maximum-minimum interval of three biological repeats with independent populations of plants. Individual biological repeats are represented with different shapes (n = 30 seedlings, 10 seedlings per biological repeat). Letters [a–b] denote groups that show statistically significant differences with other genotypes or treatments by two-way ANOVA with Tukey’s post-hoc test (P < 0.05).

When we compared the CK-666 treatment of *arpc2* with its wild-type siblings, there were similar changes in actin architecture and filament nucleation frequency (Supplemental Fig. S2 and S6). Therefore, we validated CK-666 as a small molecule inhibitor that targets the Arp2/3 complex and demonstrated its utility as an acute but reversible inhibitor for studying the Arp2/3 complex in Arabidopsis.

### The Arp2/3 complex and formins are both capable of nucleating side-branched actin filaments

Our previous work shows that profilin-bound actin monomers favor formin-mediated fast actin filament elongation at rates > 2 µm/s (Cao et al., 2016), and the data above suggest that Arp2/3-nucleated filament barbed ends elongated at intermediate rates of 1.25–1.5 µm/s (Fig. 3B; Supplemental Fig. S2B). We previously demonstrated the effectiveness of Small Molecule Inhibitor of Formin Homology 2 (SMIFH2) in Arabidopsis both in vivo and in vitro (Cao et al., 2016). The actin array organization is altered and filament nucleation frequency is significantly reduced when wild-type hypocotyl epidermal cells were treated with 25 μM SMIFH2 for 5 min (Cao et al., 2016), and similar results were obtained here (Fig. 5; Fig. 6, A and C–E; Supplemental Movie S10).

**Figure 5.**
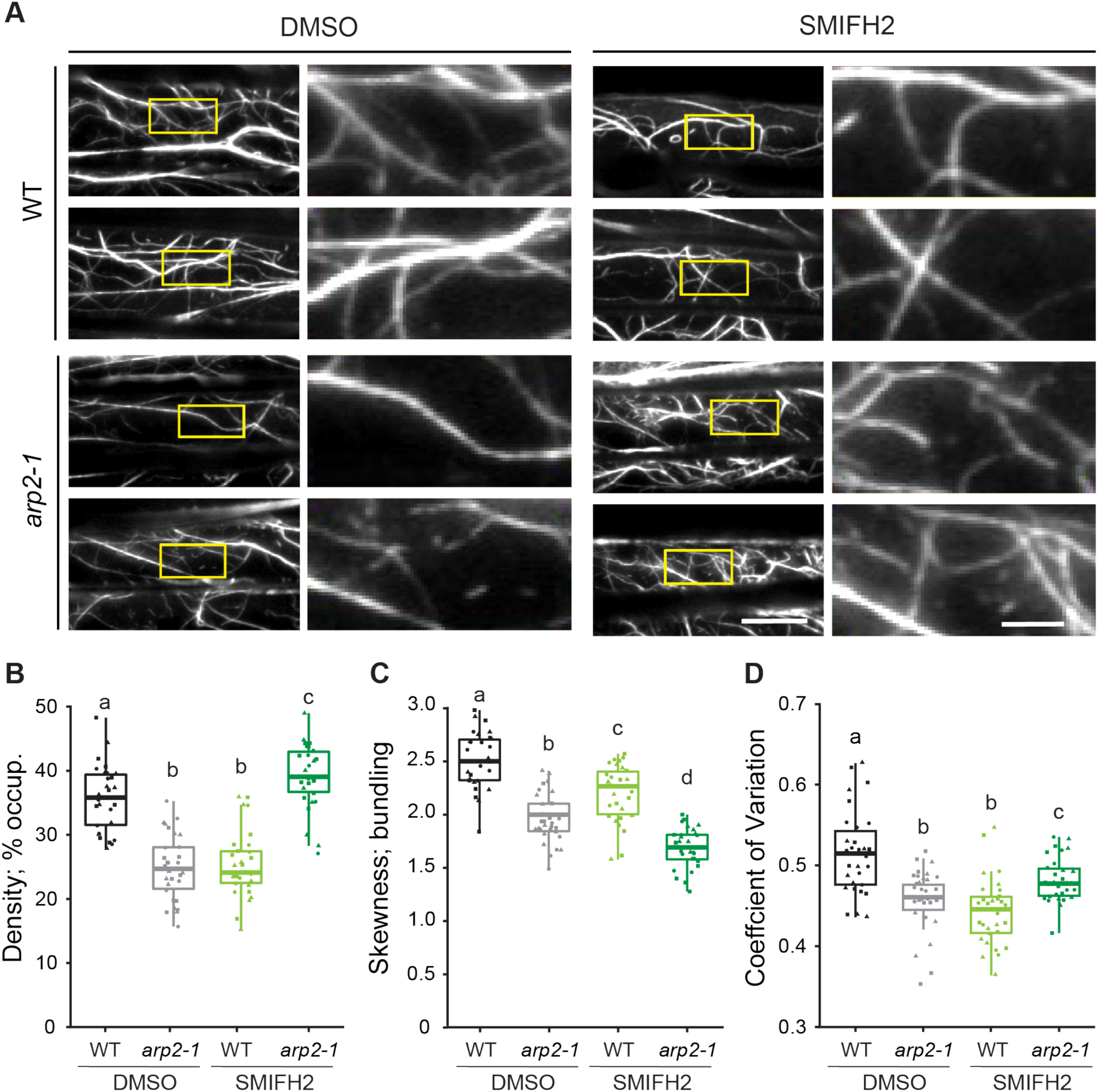
Actin filament density increases after treatment of *arp2-1* with the formin inhibitor SMIFH2. **A)** Representative images of epidermal cells from the apical region of 5-d-old etiolated hypocotyls are shown in the left column. Scale bar: 20 μm. Regions of interest (yellow boxes) were magnified and displayed in the right column. Scale bar: 5 μm. Hypocotyls were treated with 0.05% DMSO solution or 25 µM SMIFH2 for 5 min prior to imaging with VAEM. Actin filament arrays in DMSO-treated *arp2-1* and SMIFH2-treated wild-type cells appeared to be less dense and less bundled compared to DMSO-treated wild-type cells, but SMIFH2-treated *arp2-1* cells have a significantly increased actin abundance. **B–D)** Quantitative analysis of the percentage of occupancy or density of actin filament arrays **(B)**, and the extent of filament bundling as measured by skewness **(C)** and coefficient of variation **(D)** analyses. The density of actin arrays in DMSO-treated *arp2-1* and SMIFH2-treated wild-type cells was significantly decreased compared to DMSO-treated wild type; however, SMIFH2-treated *arp2-1* cells had significantly increased actin density compared to all other genotypes and treatments **(B)**. Actin arrays in *arp2-1* or SMIFH2-treated cells were significantly less bundled compared to DMSO-treated wild-type cells **(C–D)**. In box-and-whisker plots, boxes show the interquartile range (IQR) and the median, and whiskers show the maximum-minimum interval of three biological repeats with independent populations of plants. Individual biological repeats are represented with different shapes (n = 30 seedlings, 10 seedlings per biological repeat). Letters [a–d] denote groups that show statistically significant differences with other genotypes or treatments by two-way ANOVA with Tukey’s post-hoc test (P < 0.05).

**Figure 6.**
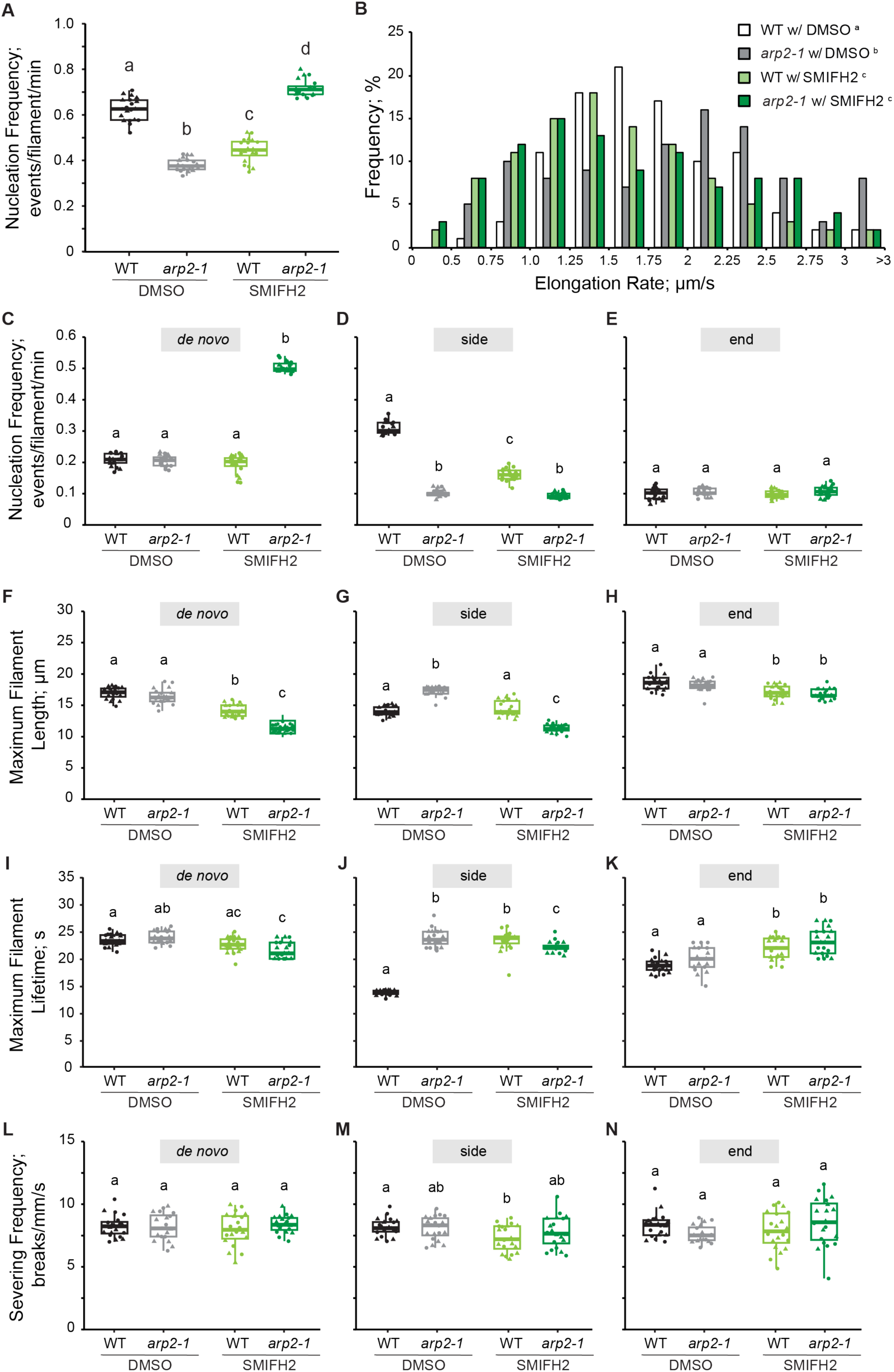
Overall and *de novo* filament nucleation increase when Arp2/3 and formin activity are simultaneously inhibited. **A and C–E)** Quantitative analysis of actin filament nucleation frequency, both overall **(A)** and by the subclass of origin **(C–E)**. The total nucleation frequency in DMSO-treated *arp2-1* and SMIFH2-treated wild-type cells was significantly reduced compared to DMSO-treated wild-type cells. However, the total nucleation frequency of SMIFH2-treated *arp2-1* cells was significantly higher than all other genotypes and treatments **(A)** and this correlated with increased *de novo* nucleation events **(C)**. In box-and-whisker plots, boxes show the interquartile range (IQR) and the median, and whiskers show the maximum-minimum interval of two biological repeats with independent populations of plants. Individual biological repeats are represented with different shapes (n = 20 seedlings, 10 seedlings per biological repeat). Letters [a–c] denote groups that show statistically significant differences with other genotypes or treatments (within the same filament nucleation subclass) by two-way ANOVA with Tukey’s post-hoc test (P < 0.05). **B)** Analysis of the population distribution of actin filament elongation rates. The elongation rate distribution of DMSO-treated wild type had a single peak at 1.25 – 1.75 μm/s, the DMSO-treated *arp2-1* had three peaks at 0.75 – 1.0 μm/s, 2.0 – 2.5 μm/s and > 3 µm/s, the SMIFH2-treated wild-type had a peak at 1.0 – 1.75 μm/s, but SMIFH2-treated *arp2-1* had peaks at 1.0 – 1.25 μm/s and 1.75 – 2.0 μm/s. n ≥ 100 single filaments from two individual biological repeats (for one biological repeat, 5 single filaments were counted in one hypocotyl from at least 10 hypocotyls per genotype or treatment). Letters [a–d] denote genotypes or treatments that show statistically significant differences with other groups by Chi-squared test, P < 0.05. **F–H)**, The average maximum length of filaments that originated *de novo* or from side-branching events in SMIFH2-treated *arp2-1* was significantly shorter than that in other cells, but filaments that originated from pre-existing ends did not show a difference between any genotype or treatment. **I–K)**, The average maximum lifetime of side-branching filaments in DMSO-treated *arp2-1*, SMIFH2-treated *arp2-1* and SMIFH2-treated wild-type cells were all significantly longer than the ones from DMSO-treated wild-type cells; however, filaments that originated *de novo* did not show any significant difference, and filaments that elongated from pre-existing ends in DMSO-treated wild-type cells were slightly shorter than SMIFH2-treated cells. **L–N)**, The severing frequency did not show any significant difference between different genotypes or treatments. In box-and-whisker plots, boxes show the interquartile range (IQR) and the median, and whiskers show the maximum-minimum interval of two biological repeats with independent populations of plants. Individual biological repeats are represented with different shapes (n = 20 seedlings, 10 seedlings per biological repeat). Letters [a–c] denote groups that show statistically significant differences with other genotypes or treatments (within the same filament nucleation subclass) by two-way ANOVA with Tukey’s post-hoc test (P < 0.05).

To test whether the homeostatic actin filament array generated by these two nucleators depends upon the Arp2/3 complex or formins, or both, we genetically and/or chemically inhibited the Arp2/3 complex and formins. Both Arp2/3-inhibited (DMSO-treated *arp2-1* or *arpc2)* or formin-inhibited (SMIFH2-treated wild-type) cells had a significant decrease in actin filament abundance as well as in the extent of filament bundling compared to DMSO-treated wild type (Fig. 5; Supplemental Fig. S7). Thus, both the Arp2/3 complex and formins are required to generate dense arrays of individual actin filaments in the homeostatic actin cortex. To test whether filament dynamics or the types and extent of filament nucleation are distinctly due to the Arp2/3 complex or formins, we examined single filament dynamics in genetically-and/or chemically-inhibited material. The nucleation frequency of both total and side-branching filaments was significantly decreased in both Arp2/3-inhibited (DMSO-treated *arp2-1*) and formin-inhibited (SMIFH2-treated wild-type) cells compared to wild-type (Fig. 6, A and C–E; Supplemental Movies S2 and S10), and the nucleation frequency of side-branched filaments in Arp2/3-inhibited cells (0.09 ± 0.01 events/filament/min) was ∼45% less than in formin-inhibited cells (0.16 ± 0.02 events/filament/min; Fig. 6D). However, the nucleation frequency due to *de novo* filament origins was not influenced when either the Arp2/3 complex or formin were suppressed (Fig. 6C). The average filament elongation rate in Arp2/3-inhibited cells (1.89 ± 0.07 μm/s) was also significantly higher than the rate in formin-inhibited cells (1.57 ± 0.06 μm/s) or in wild-type cells (1.79 ± 0.05 μm/s). In addition, the population distribution of filament elongation rates in Arp2/3-inhibited cells was significantly left-skewed (peak value at 2.0–2.5 μm/s) compared to wild type (peak value at 1.25–1.75 μm/s), whereas formin-inhibited cells lost a large portion of filaments elongating at 2.0–2.5 μm/s but the distribution had a prominent peak at 1.0–1.5 μm/s (Fig. 6B). In addition, filaments in Arp2/3-inhibited cells had a longer average length compared to formin-inhibited cells (Fig. 6, F–H), but we did not observe any significant difference in filament lifetime or severing frequency when parameters of side-branched filaments in Arp2/3-inhibited cells and formin-inhibited cells were compared (Fig. 6, I–N). Again, we found similar results in both actin architecture (Supplemental Fig. S7) and single filament dynamics (Supplemental Fig. S8) when *arpc2* and its respective wild-type line treated with DMSO solution or SMIFH2 were compared. These data indicated that both the Arp2/3 complex and formin predominantly generate side-branched actin filaments and formin-nucleated filaments elongate faster and longer than Arp2/3-nucleated ones.

To test whether SMIFH2 exhibits off-target inhibition of myosin superfamily members, as reported in Drosophila (Nishimura et al., 2021), we evaluated the influence of SMIFH2 (and CK- 666) on the activity of myosin XIK, the major myosin isoform mediating the delivery, vesicle tethering, and exocytosis of secretory vesicles in Arabidopsis cells (Zhang et al., 2021). We found that neither SMIFH2 nor CK-666 treatment reduced the speed of myosin XIK-YFP motility compared to pentabromopseudilin (PBP), an effective myosin inhibitor in plant cells (Supplemental Fig. S9, D–F) (Zhang et al., 2019). In addition, we also measured filament convolutedness and the rate of change of convolutedness of actin filaments, which are parameters describing the buckling and straightening of filaments (Staiger et al., 2009; Cai et al., 2014). Both parameters were significantly decreased after treatments with SMIFH2 or PBP, but unchanged by CK-666, indicating that SMIFH2 does indeed alter filament buckling and straightening (Supplemental Fig. S9, A–C). These data suggest that SMIFH2 may not have an off-target effect on the activity of myosins XI based on the lack of effect on the motility of YFP-XIK; however, it is remains a formal possibilitythat SMIFH2 targets myosin isoforms other than myosin XIK. Although the inhibition of filament buckling and straightening could indicate an off-target effect of SMIFH2 on myosin XI, it seems equally likely that filament buckling is caused by the processive assembly of actin filaments by Arabidopsis formins (Supplemental Fig. S9, A–C).

### Simultaneous genetic and/or chemical inhibition of Arp2/3 and formins increases spontaneous filament nucleation and overall actin filament abundance

To test whether two classes of nucleator cooperate to generate and maintain the homeostatic actin cortical array in plant epidermal cells, we explored means to simultaneously suppress the activity of both the Arp2/3 complex and formins and quantitatively evaluate the effects on actin organization and dynamics. Specifically, using chemical and genetic tools to suppress the activity of both the Arp2/3 complex and formins simultaneously, we applied SMIFH2 to *arp2-1 or arpc2* cells, treated wild-type cells with a combination of CK-666 and SMIFH2, and applied CK- 666 to the *fh1-2* mutant. Since the Arp2/3 complex and formins are, so far, the only two actin filament nucleators identified in plant cells, we predicted that simultaneously inhibiting the Arp2/3 complex and formin would markedly suppress actin filament nucleation activities and significantly reduce filament abundance. Surprisingly, in *arp2-1* cells treated with SMIFH2, the actin filament abundance was significantly increased compared to cells with normal Arp2/3 complex and formin activities or with only one nucleator inhibited (Fig. 5B). Notably, SMIFH2- treated *arp2-1* cells exhibited a marked abundance of short, single actin filaments in the cortical array (Fig. 5A). Unlike actin array density, the extent of actin filament bundling was not restored to wild-type levels when both the Arp2/3 complex and formins were simultaneously inhibited (Fig. 5, C and D). In addition to increased filament abundance, total filament nucleation frequency was significantly increased when both the Arp2/3 complex and formin activities were reduced, and a 2.5-fold enhancement of *de novo* filament origins was responsible for this increase (Fig. 6, A and C–E; Supplemental Movie S11). However, the lengths of filaments generated *de novo* or from the side-branching nucleation events were both significantly reduced in SMIFH2-treated *arp2-1* cells compared to other treatments and genotypes (Fig. 6, F–H). Similar results were obtained when *arp2-1* epidermal cells from light-grown cotyledons were treated with SMIFH2 (Supplemental Fig. S4), demonstrating that enhanced filament abundance and a 2.5-fold increase in *de novo* filament nucleation were not unique to etiolated hypocotyls. Moreover, the findings were identical when GFP-LifeAct was used as the reporter instead of GFP-fABD2 (Supplemental Fig. S3), indicating that the reporter plays little or no role in the results obtained. Finally, we also observed similar phenotypes in both actin architecture and filament dynamics when we applied SMIFH2 to *arpc2* (Supplemental Fig. S7 and S8) or simultaneously applied CK-666 and SMIFH2 to wild type (Fig. 7). These results indicate that a marked increase in *de novo* filament nucleation results when both the Arp2/3 complex and formins are inhibited in plant epidermal cells.

**Figure 7.**
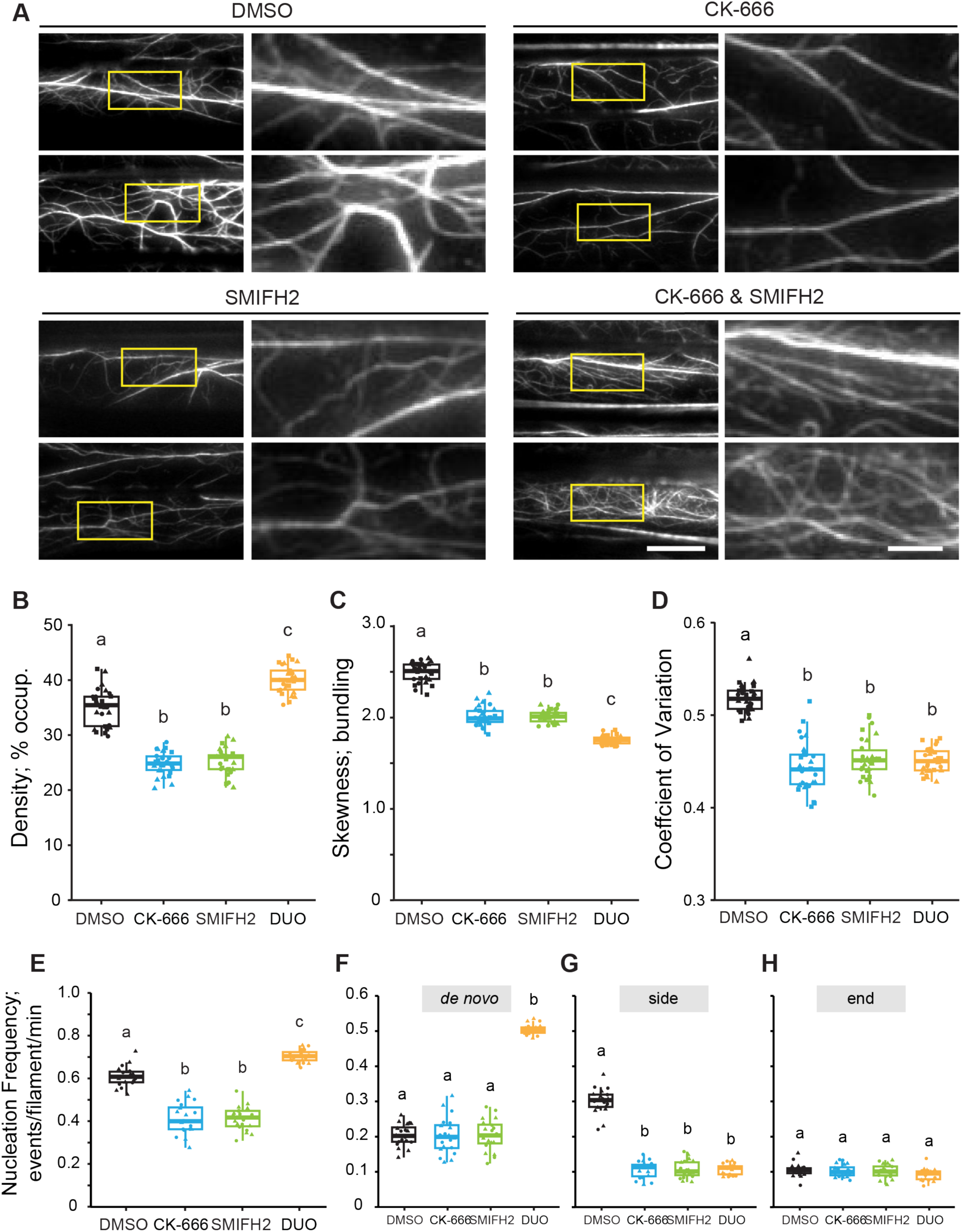
Filament abundance, total nucleation events and *de novo* filament formation increase when the Arp2/3 complex and formin activity are simultaneously reduced with chemical inhibitors. **A)** Representative images of epidermal cells from the apical region of 5-d-old etiolated hypocotyls are shown in left columns. Scale bar: 20 μm. Regions of interest (yellow boxes) were magnified and displayed in right columns. Scale bar: 5 μm. Hypocotyls were treated with 0.05% DMSO solution, 10 µM CK-666, 25 µM SMIFH2 or both inhibitors for 5 min prior to imaging with VAEM. Actin filament arrays in CK-666-treated cells and SMIFH2-treated cells appeared to be less dense and less bundled compared to mock-treated cells. However, dual treatment markedly increased actin filament abundance. **B–D)** Quantitative analysis of actin filament density **(B)** and extent of bundling by skewness **(C)** and coefficient of variation **(D)** analyses. The density of actin arrays in CK-666-treated and SMIFH2-treated cells was decreased compared to mock-treated cells; however, dual-treated (DUO) cells had significantly increased actin density compared to all other treatments. Actin arrays in CK-666- or dual-treated cells were significantly less bundled than in mock-treated cells. In box-and-whisker plots, boxes show the interquartile range (IQR) and the median, and whiskers show the maximum-minimum interval of three biological repeats with independent populations of plants. Individual biological repeats are represented with different shapes (n = 30 seedlings, 10 seedlings per biological repeat). Letters [a–c] denote groups that show statistically significant differences with other genotypes or treatments by two-way ANOVA with Tukey’s post-hoc test (P < 0.05). **E–H)** Quantitative analysis of actin filament nucleation frequency, both overall **(E)** and by subclass of origin **(F– H)**. The total nucleation frequency in mock-treated cells was higher than CK-666- treated and SMIFH2-treated cells. However, the total nucleation frequency of dual-treated (DUO) cells was significantly higher than all other treatments and this correlated with increased *de novo* nucleation events. In box-and-whisker plots, boxes show the interquartile range (IQR) and the median, and whiskers show the maximum-minimum interval of two biological repeats with independent populations of plants. Individual biological repeats are represented with different shapes (n = 20 seedlings, 10 seedlings per biological repeat). Letters [a–c] denote groups that show statistically significant differences with other genotypes or treatments by two-way ANOVA with Tukey’s post-hoc test (P < 0.05).

Beside the possible inhibition of myosin XI, another potential issue with using SMIFH2 as a formin inhibitor on plant cells is that Arabidopsis has 21 FORMIN homologs, and it remains unclear whether SMIFH2 can inhibit all of them. To investigate the efficacy of SMIFH2 for suppressing formin activity, as well as its ability to phenocopy the effects of plant formin mutants, we prepared a homozygous *AtFORMIN1* mutant line, *fh1-2*, expressing GFP-fABD2. AtFORMIN1 is a major housekeeping formin homolog in Arabidopsis vegetative tissues (Cvrčková et al., 2000; Blanchoin and Staiger, 2010; Rosero et al., 2013; Rosero et al., 2016). Previously, we demonstrated that SMIFH2 inhibits the nucleation and assembly activity of recombinant FH2 domain from AtFORMIN1 in vitro (Cao et al., 2016). Here, we found that the density of actin filament arrays was significantly reduced but the extent of filament bundling was higher in DMSO-treated *fh1-2* cells compared to DMSO-treated wild-type (Fig. 8, A–D). The nucleation frequency, specifically the side-branched filament subclass, was also significantly decreased in DMSO-treated *fh1-2* cells compared to DMSO-treated wild-type (Fig. 8, E–H). SMIFH2 treatment caused an additional reduction in both actin filament abundance and nucleation activity in *fh1-2* cells (Fig. 8). These results suggest that SMIFH2 targets more formin homologs than just AtFORMIN1; however, we cannot rule out the possibility that SMIFH2 does not inhibit all formins. Consistent with previous biochemical results (Michelot et al., 2007), these findings provide genetic evidence that AtFORMIN1 is a filament nucleator that generates new daughter filaments from the side of a mother filament or bundle.

**Figure 8.**
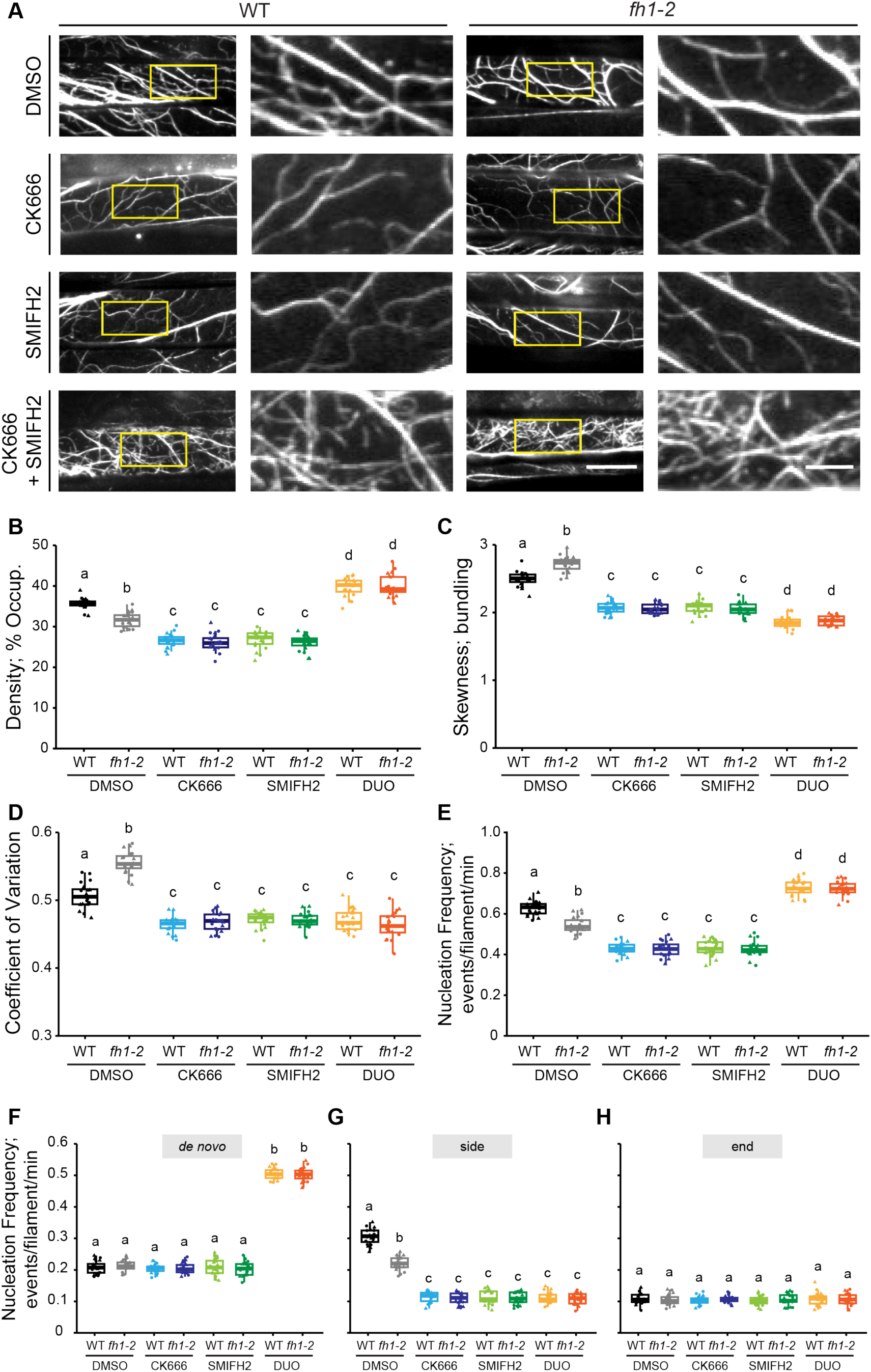
SMIFH2 treatment results in further reduction of both actin density and actin filament nucleation frequency in the *fh1-2* mutant. **A)** Representative images of epidermal cells from the apical region of 5-d-old etiolated hypocotyls are shown in the left columns. Scale bar: 20 μm. Regions of interest (yellow boxes) were magnified and displayed in the right columns. Scale bar: 5 μm. Hypocotyls were treated with 0.05% DMSO solution, 10 µM CK-666, 25 µM SMIFH2 or both inhibitors for 5 min prior to imaging. Actin filament arrays in either DMSO-treated or single-inhibitor-treated *fh1-2* cells all appeared to be less dense compared to DMSO-treated wild-type cells, but both dual-inhibitor-treated wild-type and *fh1-2* cells appeared to have more dense actin arrays compared to DMSO-treated wild-type cells. **B–D)** Quantitative analysis of the percentage of occupancy or density of actin filament arrays **(B)** and the extent of filament bundling as measured by skewness **(C)** and coefficient of variance **(D)** analyses. The density of actin arrays in DMSO-treated *fh1-2* cells was significantly decreased but the extent of filament bundling was increased compared to DMSO-treated wild-type cells. Treatment with either CK-666 or SMIFH2 caused an additional decrease in actin density in *fh1-2* cells and filaments were less bundled compared to DMSO-treated wild-type. However, simultaneous treatment with both inhibitors (DUO) significantly increased actin array density in both wild-type and *fh1-2* cells. In box-and-whisker plots, boxes show the interquartile range (IQR) and the median, and whiskers show the maximum-minimum interval of two biological repeats with independent populations of plants. Individual biological repeats are represented with different shapes (n = 20 seedlings, 10 seedlings per biological repeat). Letters [a–d] denote groups that show statistically significant differences with other genotypes or treatments by two-way ANOVA with Tukey’s post-hoc test (P < 0.05). **E–H)** Quantitative analysis of actin filament nucleation frequency, both overall **(E)** and by subclass of origin **(F–H)**. The total nucleation frequency in DMSO-treated *fh1-2* cells was significantly reduced compared to DMSO-treated wild-type cells, and either CK-666 or SMIFH2 treatment caused an additional decrease in overall filament nucleation in *fh1-2* cells **(E)**. However, the total nucleation frequency of dual-inhibitor-treated *fh1-2* or wild-type cells was significantly higher than all other genotypes and treatments **(E)** and this correlated with increased *de novo* nucleation events **(F)**. In box-and-whisker plots, boxes show the interquartile range (IQR) and the median, and whiskers show the maximum-minimum interval of two biological repeats with independent populations of plants. Individual biological repeats are represented with different shapes (n = 20 seedlings, 10 seedlings per biological repeat). Letters [a–d] denote groups that show statistically significant differences with other genotypes or treatments by two-way ANOVA with Tukey’s post-hoc test (P < 0.05).

To further test whether loss of a single formin homolog could recapitulate the effects of SMIFH2, we treated *fh1-2* with CK-666 or a combination of CK-666 and SMIFH2, as shown in Figure 8. Treatment of *fh1-2* with CK-666 led to a further significant reduction in filament density (Fig. 8B) as well as side-branched nucleation frequency (Fig. 8G), but no change in *de novo* nucleation events (Fig. 8F), compared to DMSO-treated *fh1-2*. This does not resemble the dual inhibition of wild-type with SMIFH2 and CK-666 (Fig. 7), suggesting that loss of a single major formin is not responsible for the phenotype. Further, we found that identical to wild-type cells simultaneously treated with SMIFH2 and CK-666, *fh1-2* responds to dual inhibition with significantly increased filament abundance (Fig. 8B) as well as enhanced total (Fig. 8E) and *de novo* nucleation frequency (Fig. 8F). These results indicate that loss of AtFH1 does not play a role in the rapid response to inhibition of two classes of nucleator in epidermal cells resulting in enhanced filament formation through *de novo* nucleation.

### PROFILIN1 does not play a role in the enhanced *de novo* nucleation in response to simultaneous inhibition of two classes of nucleator

Profilins are actin monomer-binding proteins that suppress spontaneous nucleation, prevent subunit addition onto filament pointed ends (Cao et al., 2016; Sun et al., 2018), and are reportedly present at up to a 3-fold molar excess to total actin protein in several plant tissues (Chaudhry et al., 2007). To test whether the enhanced *de novo* actin filament nucleation, when the Arp2/3 complex and formins are both inhibited, was due to spontaneous nucleation, we introduced a PROFILIN1 (PRF1) mutant, *prf1-2*, to suppress one of the major profilin homologs in Arabidopsis vegetative tissues (Sun et al., 2018; Qiao et al., 2019). Consistent with our previous findings (Cao et al., 2016), *prf1-2* had significantly reduced actin filament abundance (Fig. 9, A and B), decreased total and side-branched nucleation (Fig. 9, E and G), but modestly elevated *de novo* nucleation frequency (Fig. 9F) compared to mock-treated wild type. If the enhanced *de novo* nucleation frequency following simultaneous inhibition of both nucleators is due to the spontaneous nucleation resulting from increased free actin monomers, then we expect that *prf1-2* would be less responsive to these conditions as it presumably already has an elevated free monomer concentration. We found that the triple inhibition of Arp2/3, formins and PRF1 caused a significantly increased actin density (Fig. 9, A and B), as well as enhanced total and *de novo* nucleation frequency (Fig. 9, E–H), all of which were slightly elevated compared with dual-inhibitor treated wild type. However, either CK-666 or SMIFH2 single inhibitor treatment on *prf1-2* did not cause a 2.5-fold higher *de novo* nucleation frequency (Fig. 9E) as observed in dual-inhibitor treated *prf1-2* cells. These results suggest that a change in free actin monomer concentration or the function of PRF1 are not the cause of the elevated *de novo* nucleation when both the Arp2/3 complex and formins are inhibited.

**Figure 9.**
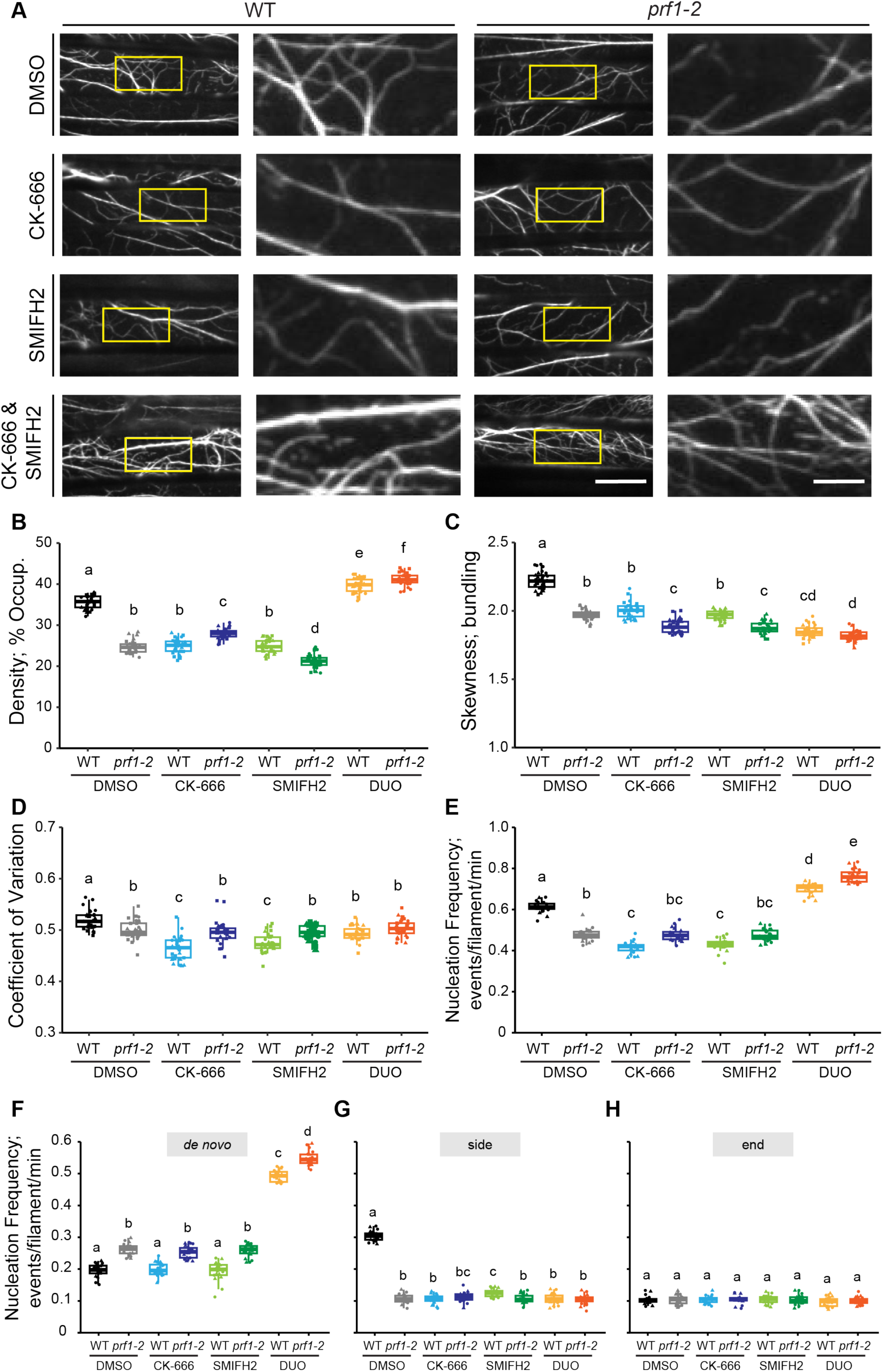
Increased filament abundance, total nucleation events and *de novo* filament formation following simultaneous CK-666 and SMIFH2 treatment are further enhanced in the *prf1-2* mutant. **A)** Representative images of epidermal cells from the apical region of 5-d-old etiolated hypocotyls are shown in the left columns. Scale bar: 20 μm. Regions of interest (yellow boxes) were magnified and displayed in the right columns. Scale bar: 5 μm. Hypocotyls were treated with 0.05% DMSO solution, 10 µM CK-666, 25 µM SMIFH2 or both inhibitors for 5 min prior to imaging with VAEM. Actin filament arrays in either DMSO-treated or single-inhibitor-treated *prf1-2* cells all appeared to be less dense and less bundled compared to DMSO-treated wild-type cells, but both dual-inhibitor-treated wild-type and *prf1-2* cells appeared to have more dense actin arrays compared to DMSO-treated wild-type cells. **B–D)** Quantitative analysis of actin filament density **(B)** and extent of bundling by skewness **(C)** and coefficient of variation **(D)** analyses. The density of actin arrays in either DMSO-treated or single-inhibitor-treated *prf1-2* cells was significantly decreased compared to DMSO-treated wild-type cells; however, the treatment with both inhibitors (DUO) significantly increased the actin density in both wild-type and *prf1-2* cells. Actin arrays were less bundled when either Arp2/3, formins or PRF1 were inhibited compared to DMSO-wild-type cells. In box-and-whisker plots, boxes show the interquartile range (IQR) and the median, and whiskers show the maximum-minimum interval of three biological repeats with independent populations of plants. Individual biological repeats are represented with different shapes (n = 30 seedlings, 10 seedlings per biological repeat). Letters [a–f] denote groups that show statistically significant differences with other genotypes or treatments by two-way ANOVA with Tukey’s post-hoc test (P < 0.05). **E–H)** Quantitative analysis of actin filament nucleation frequency, both overall **(E)** and by subclass of origin **(F–H)**. The total nucleation frequency in DMSO-treated, CK-666-treated and SMIFH2-treated *prf1-2* cells was significantly reduced compared to DMSO-treated wild-type cells. However, the total nucleation frequency of dual-inhibitor-treated (DUO) *prf1-2* **(E)** was significantly higher than all other genotypes and treatments and this corresponded to a significant increase in *de novo* nucleation events **(F)**. In box-and-whisker plots, boxes show the interquartile range (IQR) and the median, and whiskers show the maximum-minimum interval of two biological repeats with independent populations of plants. Individual biological repeats are represented with different shapes (n = 20 seedlings, 10 seedlings per biological repeat). Letters [a–e] denote groups that show statistically significant differences with other genotypes or treatments by two-way ANOVA with Tukey’s post-hoc test (P < 0.05).

## Discussion

In this study, we utilized a combination of genetic mutations or small molecule inhibitors along with high spatiotemporal resolution fluorescence microscopy to demonstrate, for the first time, that the Arp2/3 complex nucleates side-branched actin filaments in living Arabidopsis epidermal cells. We found that the Arp2/3 complex mutants, *arp2-1* and *arpc2*, had a dramatically reduced overall abundance of actin filaments in the cortical array, a reduction in the extent of filament bundles, and a significantly decreased frequency of side-branched nucleation events. In addition, acute treatment of wild-type plants with CK-666 phenocopied the actin-based defects observed in *arp2/3* mutants, whereas applying CK-666 to *arp2/3* mutants did not have any additional effect on overall actin structure or filament dynamics. Thus, we confirmed that these actin-based defects correlated with the loss of functional Arp2/3 complex and established that CK-666 is an effective tool for inhibition of the Arp2/3 complex in plant cells. Through experiments comparing genetic and chemical inhibition of the Arp2/3 complex or formins, we observed that both proteins were capable of nucleating side-branched actin filaments, but that Arp2/3-nucleated filaments grew slower and were shorter than formin-nucleated ones. Surprisingly, simultaneous inhibition of both the Arp2/3 complex and formins led to an increase in actin filament abundance and dramatically promoted *de novo* nucleation events. By examining the consequences of the loss of a major housekeeping profilin isoform, PRF1, we rule out the role of this monomer-binding protein in the enhanced *de novo* nucleation following simultaneous inhibition of two classes of nucleator. These observations indicate that the regulatory mechanisms for actin filament nucleation in the homeostatic cortical array of plant cells may be unique and reveal a default response that maintains filament abundance and dynamics when both nucleators are inactivated.

### The Arp2/3 complex is a major nucleator of side-branched actin filaments in *Arabidopsis* epidermal cells

Biochemical studies demonstrate that the Arp2/3 complex and formins use different molecular mechanisms to overcome the rate-limiting step for filament formation, and they typically generate different types of actin arrays. After creating an actin nucleus, the Arp2/3 complex remains attached to the pointed end of the new filament so that the barbed end is free for the addition of actin monomers, profilin-actin, or capping protein (Mullins et al., 1998; Amann and Pollard, 2001; Fujiwara et al., 2002; Ichetovkin et al., 2002; Rouiller et al., 2008). In contrast, many formins remain associated at the filament barbed end after the nucleation, and processively facilitate filament elongation through the addition of profilin-actin. This allows formin-associated filaments to grow much faster and longer compared to filaments with free barbed ends that are typically bound by heterodimeric capping protein (CP) soon after initiation (Kovar and Pollard, 2004; Romero et al., 2004; Kovar, 2006). Based on these observations, the conventional model of actin nucleation posits that formins generate long, unbranched filaments with a fast filament elongation rate, whereas the Arp2/3 complex nucleates short, branched actin networks with a slower growth rate. The Arp2/3 complex is accepted as a major nucleator of side-branched actin filaments in yeast and animal cells (Mullins et al., 1998; Amann and Pollard, 2001; Fujiwara et al., 2002; Ichetovkin et al., 2002; Rouiller et al., 2008). Even though the Arp2/3 complex is also shown to be critical for plant cells to maintain their homeostatic actin organization and is required for a wide variety of cellular activities, little is known about the exact function of plant Arp2/3 complex in terms of its contribution to actin filament dynamics.

In this study, we characterized the function of the Arp2/3 complex in coordinating the organization of the cortical actin array as well as single actin filament dynamics in living Arabidopsis cells. Epidermal cells of etiolated hypocotyls, as well as light-grown cotyledons, with genetically-or chemically-inhibited Arp2/3 complex showed a quantifiable decrease in both actin filament abundance and extent of filament bundling, and, more importantly, there was a significantly reduced nucleation frequency of side-branched actin filaments. Branched filament formation represents more than half of all nucleation events in the cortical actin array of hypocotyl or cotyledon epidermal cells, and these were reduced by 50–70% when the Arp2/3 complex was inhibited chemically or genetically. These results suggest that the Arp2/3 complex is a major nucleator of side-branched actin filaments in Arabidopsis epidermal cells. Our data also provide a compelling live-cell view of the formation of branched actin filaments by active Arp2/3 complex at single filament resolution. Furthermore, our results reveal that there are at least two populations of filaments with different dynamic properties, likely due to different filament growth mechanisms and/or the availability of filament barbed ends for monomer addition, that contribute to the organization of the homeostatic actin cortical array in epidermal cells. By examining filament elongation rate frequencies when formin activity was suppressed, or when both the Arp2/3 complex and formins were inhibited, filaments with elongation rates of 1–1.25 µm/s predominated. We expect this represents the population of filaments with free barbed ends extending by monomer addition and is consistent with the in-vitro mechanism of branched filament nucleation by the Arp2/3 complex.

### The Arp2/3 complex and formins both mediate side-branched filament nucleation but generate filaments with unique dynamic properties

Nucleation is the rate-limiting step for actin filament formation in vitro and depends on the availability of actin monomers and is suppressed by profilin (Pollard et al., 2000; Sept and McCammon, 2001; Michelot et al., 2005). Previous results with the fission and budding yeasts, *Schizosaccharomyces pombe* and *Saccharomyces cerevisiae*, as well as mammalian cells reveal a competition between the Arp2/3 complex and formin for a limited pool of actin monomers; consequently, formin-mediated long filament bundles dominate when the Arp2/3 complex is inhibited, whereas an increased abundance of actin patches created by the Arp2/3 complex are prominent when formin is down-regulated (Hotulainen and Lappalainen, 2006; Burke et al., 2014; Lomakin et al., 2015; Suarez et al., 2015; Fritzsche et al., 2016; Davidson et al., 2018; Antkowiak et al., 2019; Chan et al., 2019; Kadzik et al., 2020). Plant cells, by contrast, appear to show cooperation between these two nucleators rather than a competition to build distinct actin arrays. The cortical cytoplasm of epidermal cells contains a dynamic actin array that is rather disordered and comprises comingled actin filament bundles and individual actin filaments (Staiger et al., 2009; Smertenko et al., 2010). Evidence for actin patches or dense dendritic actin networks is limited to certain cell types, such as guard mother cells (Facette et al., 2015), the apex of trichomes (Yanagisawa et al., 2015), or at focal sites elicited by pathogen attack (Hardham et al., 2007; Qin et al., 2021). Our results revealed that the density of actin filaments in the cortical array, the extent of filament bundling, as well as filament nucleation frequency all decreased when either the Arp2/3 complex or formins were inhibited, suggesting that the Arp2/3 complex or formins alone are not able to maintain the homeostasis of the cortical actin arrays when the other nucleator is not functional. These results indicate that plant cells may have a different actin regulatory mechanism for constructing actin filament arrays compared to yeast or some animal cells. Our results are consistent with a previous report showing partial cooperation between plant filament nucleation mechanisms in Arabidopsis cotyledon epidermal cells, however, the consequences of losing either class of nucleator were markedly different (Cifrová et al., 2020). That study showed a minor increase in actin filament density in either the formin *fh1* mutant or the *arpc5* mutant (Cifrová et al., 2020), rather than the prominent decreases observed here. As demonstrated here, these differences are not due to the use of different actin filament reporters (LifeAct versus fABD2), different developmental state and organ for examining epidermal cells (cotyledons versus dark-grown hypocotyls), or the result of compensation from genetic loss of one formin isoform versus acute treatment with a chemical inhibitor.

Our results showed that both the Arp2/3 complex and formins, including AtFORMIN1, are responsible for generating side-branched filaments because the inhibition of either one led to a significant reduction in side-branched filament nucleation and did not promote the other two subclasses of filament nucleation. These results also suggest that the Arp2/3 complex and formins are not competing for actin monomers for filament nucleation. Based on the total actin concentration, the ratio of F-actin to total actin, and the ratio of profilin to total actin in other plant tissues, the estimated actin monomer pool in Arabidopsis hypocotyl cells is relatively large and most monomers are likely to be in a profilin-actin complex (Chaudhry et al., 2007; (Staiger et al., 2010); therefore, the Arp2/3 complex and formins may not have to compete for a limited supply of actin monomers in plant cells. Here, if we assume that 1.25 µm/s is the rate of addition of monomers to free barbed ends, that the association rate constant for plant actin is similar to ATP-loaded rabbit skeletal muscle α-actin (k_+_ = 11.6 µM^-1^ s^-1^) and that a micron of actin filament comprises 370 subunits (Pollard et al., 2000), then we estimate the available monomer pool to be 40 µM in hypocotyl epidermal cells using the equation: [G-actin] = rate/k_+_. In addition, even though both the Arp2/3 complex and formins generate side-branched actin filaments, these filaments have markedly different properties. We found that formin-nucleated filaments grew significantly faster at rates of 2–2.25 µm/s, with another small subpopulation growing at rates > 3 µm/s. The different properties of filament barbed ends as well as a large pool of monomers, perhaps buffered with profilin, may facilitate the fast-growing filaments nucleated by side-branching as previously found in Arabidopsis cells (Cao et al., 2016), and is consistent with previous in vitro results (Vavylonis et al., 2006; Michelot et al., 2013; Suarez et al., 2015; Zhang et al., 2016; Funk et al., 2019).

If using the conventional model for how formins generate new actin filaments, we would predict that plant processive formins not only generate growing filaments from the side of a mother filament but also remain attached to the barbed end of the new filament to facilitate the addition of profilin-actin to the growing ends, resulting in faster and longer side-branched filaments compared to Arp2/3-nucleated ones. However, previous in vitro data showed that the AtFORMIN1 nucleates actin filaments but remains associated at the pointed end and does not compete with capping protein for filament barbed ends, indicating it is non-processive and associates with the side of filaments after nucleation (Michelot et al., 2006). Further, SMIFH2 application in our studies reduced nucleation frequency but did not reduce the filament length of side-branched filaments compared to the mock treatment. Both in vitro and in vivo results suggest that non-processive formins in plant cells may have a weak interaction with filament barbed ends so that these formins move away from the end of the filament to its side to generate a new side-branched filament instead of occupying the barbed end and facilitating filament elongation. However, another in vitro analysis showed that the processive formin, AtFORMIN14, promotes *de novo* filament growth via the association with profilin (Zhang et al., 2016), and there are 21 FORMIN homologs in *Arabidopsis* (Cvrčková et al., 2004; Blanchoin and Michelot, 2012; Cvrčková, 2013). All these results suggest that plant formins that nucleate side-branched filaments may have distinct molecular properties compared to formins that nucleate *de novo* filament formation or formins found in yeast or animal cells. With the exception of enhanced bundling, the loss of a single formin isoform, *fh1-2*, recapitulates but is less dramatic than the effects of SMIFH2 treatment of epidermal cells supporting the conclusion that multiple formin isoforms are likely inhibited by this small molecule.

Since SMIFH2 shows off-target inhibition of members of the myosin superfamily (Nishimura et al., 2021), we were compelled to critically evaluate the effects of SMIFH2 treatment on plant cells. Although our data showed that SMIFH2 treatment did not influence the motility of myosin XIK, it did significantly reduce filament convolutedness and the rate of change of convolutedness of actin filaments, suggesting that SMIFH2 alters filament buckling and straightening. However, the change in filament buckling could be the consequence of different filament growth rates and processive elongation by formins (Staiger et al., 2009; Cai et al., 2014; Cao et al., 2016). Even though a recent in vitro study reveals that SMIFH2 is a pan-inhibitor for all human formins (Orman et al., 2022), we should not exclude the possibility that SMIFH2 can only inhibit a subset of plant formin homologs. Further genetic and biochemical analyses should be designed to dissect the interaction between SMIFH2 and different formin homologs to determine the specificity of this chemical inhibitor on plant formins.

### Simultaneous inhibition of the Arp2/3 complex and formins promotes *de novo* nucleation events

When we combined genetic and chemical methods to downregulate the Arp2/3 complex and formins simultaneously, surprisingly, the frequency of actin filament nucleation was significantly enhanced by up to 2.5-fold, with *de novo* nucleation events appearing to be the exclusive contributor to this enhancement. The Arp2/3 complex and formins are, to date, the only two *bona fide* actin filament nucleators in plant cells (Blanchoin and Staiger, 2010; Vaškovičová et al., 2013; Yanagisawa et al., 2013). Theoretically, when both the Arp2/3 complex and formins are inhibited, the concentration of free actin monomers is predicted to increase but the ratio of F-to G-actin and the proportion of profilin-actin in the monomer pool should both decrease. It is possible that these changes in the actin monomer pool could increase the nucleation rate of spontaneous filament formation thereby enhancing the total frequency of *de novo* nucleation events. In support of this model, disruption of AtPRF1 led to a modest increase in *de novo* nucleation events and a decrease in side-branched events (Cao et al., 2016). However, the *prf1-2* mutant exhibits a 2.5-fold increase in *de novo* nucleation frequency, just like wild-type epidermal cells, when both nucleators are inhibited with chemicals. Perhaps, as suggested by Sun et al. (2018), another profilin isoform, PRF3, is the major regulator of spontaneous filament nucleation and sequesters actin monomers that are poorly utilized by formins. The acute chemical inhibition of nucleation machinery could lead to rapid post-translational modification and inactivation of PRF3, thereby facilitating spontaneous filament formation from free monomers. This hypothesis needs to be tested through future experiments.

In conclusion, our results provide direct evidence to demonstrate that the Arp2/3 complex nucleates side-branched actin filaments in living plant cells. These studies also confirm that some formin isoforms, including AtFORMIN1, generate side-branched actin filaments in plant cells, which is a unique actin filament nucleation mechanism compared to yeast or animal cells. Moreover, we find that Arp2/3- and formin-nucleated filaments have distinct dynamic properties, which can be applied to future studies about how cells coordinate these two nucleators to regulate the actin structure and dynamics for different cellular activities. The simultaneous inhibition of both the Arp2/3 complex and formin surprisingly caused an enhanced *de novo* filament nucleation, which raises the possibility for a unique actin nucleation mechanism to generate dynamic cortical actin arrays in plant cells.

## Materials and methods

### Plant materials and growth conditions

All Arabidopsis (*Arabidopsis thaliana*) plants in this study were in Columbia-0 (Col-0) background. The *ARP2* T-DNA insertion line, *arp2-1* (SALK_003448), the *PRF1* T-DNA insertion line, *prf1-2* (SALK_057718; Cao et al., 2016), and the *AtFORMIN1* T-DNA insertion line, *fh1-2* (SALK_009693; Rosero et al., 2013) were obtained from the Arabidopsis Biological Resource Center (Ohio State University). The *ARPC2* point mutation line, *arpc2* (G1217A), was kindly provided by Dan Szymanski (Purdue University). Mutant lines, *arp2-1, arpc2* and *prf1-2*, were crossed to wild-type (Col-0) expressing the GFP-fABD2 reporter (Sheahan et al., 2004; Staiger et al., 2009) and homozygous mutants and corresponding wild-type siblings were recovered from F2 populations. For the *fh1-2* line, homozygous plants were transformed with a *35S::GFP-fABD2* construct using the floral dip method and seeds were screened with kanamycin (Clough and Bent, 1998; Sheahan et al., 2004). For lines carry GFP-LifeAct, both Col-0 and *arp2-1* mutant were transformed with a *UBQ::GFP-LifeAct* construct (kindly provided by Weibing Yang from The Sainsbury Laboratory, University of Cambridge, Cambridge, UK) using the floral dip method and transformants selected with BASTA (Clough and Bent, 1998).

Seeds were surface sterilized and stratified at 4°C for 3 days on plates comprising half-strength Murashige and Skoog (MS) medium supplemented with 1% (w/v) sucrose and 1% (w/v) agar. For dark-grown hypocotyl growth, seedlings were grown in continuous darkness after exposing the plates to light for 4 h. For light-grown root and cotyledon growth, seeds were grown vertically on plates comprising half-strength MS medium supplemented with 0% (w/v) sucrose and 0.6% (w/v) agar under long-day conditions (16 h light/8h dark) at 21°C.

### Drug treatments

For short-term live cell treatments, individual seedlings were treated for 5 min in inhibitor solution in 6-well plates in the dark prior to each imaging session. Hypocotyls were mounted in the inhibitor solution and imaged immediately. CK-666 (Sigma-Aldrich; Cat # 182515; St. Louis, MO, USA), SMIFH2 (Sigma-Aldrich; Cat # 344092) and PBP (Adipogen; Cat # BVT-0441) were dissolved in DMSO to prepare a 5 mM stock solution.

### Live cell imaging

For both snapshot and time-lapse image acquisitions in dark grown hypocotyls, epidermal cells from the apical region of 5-d-old dark-grown hypocotyls were used. In some experiments, the lipophilic dye, FM4-64 (Invitrogen; Cat # T3166; Carlsbad, CA, USA), was dissolved in DMSO and used at 20 mM to label the plasma membrane. All images and movies were collected using variable-angle epifluorescence microscopy (VAEM) with an Olympus TIRF objective (60X, 1.45 numerical aperture) using SlideBook software (version 5.5; Intelligent Imaging Innovations) as described previously (Staiger et al., 2009; Henty et al., 2011; Li et al., 2012; Cai et al., 2014; Cao et al., 2016).

For experiments with cotyledons, adaxial epidermal cells from 5-d-old light-grown cotyledons were used. An Olympus IX-83 microscope mounted with a spinning-disc confocal unit (Yokogawa CSUX1-A1) and an Andor iXon Ultra 897BV EMCCD camera were used to acquire snapshot and time-lapse images with an Olympus UPlanSApo oil objective (100X, 1.45 numerical aperture). GFP fluorescence was excited with a 488nm laser line and emission collected through the 525/30 nm filter to visualize actin filaments. All images were collected with MetaMorph software (version 7.8.8.0) as described previously (Cao et al., 2022).

A double-blinded experimental design was used for all image collection, processing and quantitative analyses.

### Image processing and quantitative analysis of actin cytoskeleton architecture and single actin filament dynamics

Images and movies were processed with Fiji is Just Image J (Schindelin et al., 2012). All images used for actin architecture analysis were collected with a fixed exposure time, laser power, and camera gain setting.

To quantify actin architecture, three parameters, *density*, *skewness* and *coefficient of variation (CV)*, were employed in this study; density measures the percentage of occupancy of GFP signal in an image and skewness and CV measure the extent of actin filament bundling(Higaki et al., 2010; Ueda et al., 2010; Henty et al., 2011; Li et al., 2012; Cai et al., 2014; Cao et al., 2016; Arieti and Staiger, 2020). Micrographs were analyzed in Fiji Is Just Image J using the methods described previously (Higaki et al., 2010; Henty et al., 2011; Li et al., 2012; Higaki et al., 2020).

To compare single actin filament parameters between genotype or drug treatment, double-blind experiments (data collection and analysis) were performed. Maximum filament length and lifetime, filament severing frequency, elongation rates, filament convolutedness and the rate of change of convolutedness were measured as described previously (Staiger et al., 2009; Henty et al., 2011; Li et al., 2012; Cai et al., 2014; Cao et al., 2016). For nucleation frequency analysis, a 400 µm^2^ ROI was randomly selected from movies of epidermal cells at the apical region of the hypocotyl, and all observable nucleation events were counted during a time period of 100 s, according to Cao et al., 2016. To account for differences in filament density in genotypes or drug treatments, the nucleation frequency was normalized against filament numbers in each ROI.

For actin architecture analysis in both hypocotyl and cotyledon epidermal cells, more than 150 ROIs were selected from 50 images of cells collected from at least 10 individual seedlings per genotype or treatment. For nucleation frequency analysis, more than 300 events from at least 10 individual seedlings were observed per genotype or treatment. For single actin filament dynamics analysis, at least 5 newly-appeared filaments were tracked from their first appearance to their complete disappearance in each hypocotyl, and more than 50 filaments from at least 10 hypocotyls were measured per genotype or treatment. For all statistical comparisons and plotting, the average value from a single hypocotyl was used as one data point. All experiments were conducted with at least 2 independent biological repeats to make conclusions in each figure.

### Statistical analyses

Two-way ANOVA with Tukey’s post-hoc tests were performed in SPSS (version 25) to determine the significance among different genotypes/treatments. Chi-square tests were performed for statistically comparing parametric distributions and P values were calculated in Excel 15.32. Any difference with a P value less than 0.05 was considered significantly different. Detailed statistical analysis data are shown in Supplemental Data Set 1.

## Supplemental data

The following materials are available in the online version of this article.

**Supplemental Table S1.** Single actin filament dynamics in wild type (*ARPC*2) and *arpc2* mutant with or without CK-666.

**Supplemental Figure S1.** Disruption of the Arp2/3 complex perturbs epidermal cell morphology and inhibits normal plant growth.

**Supplemental Figure S2.** Actin filament nucleation frequency is decreased in *arpc2* and after treatment of wild type with CK-666.

**Supplemental Figure S3.** Two different actin cytoskeleton markers, GFP-fABD2 and GFP-LifeAct, report similar changes in actin array organization and nucleation events regardless of the genotype or treatment.

**Supplemental Figure S4.** Cotyledon epidermal cells show similar phenotypes of reduced actin filament abundance and decreased nucleation frequency in *arp2-1* mutants expressing either GFP-fABD2 or GFP-LifeAct cytoskeletal reporters.

**Supplemental Figure S5.** CK-666 is an acute and reversible chemical inhibitor of the Arp2/3 complex in plant cells.

**Supplemental Figure S6.** Actin array architecture is perturbed in *arpc2* and the reduced filament density and bundling phenotypes are recapitulated by treatment with CK-666.

**Supplemental Figure S7.** Actin filament density increases after treatment of *arpc2* with the formin inhibitor SMIFH2.

**Supplemental Figure S8.** Overall and *de novo* filament nucleation increase when ARPC2 and formin activity are simultaneously inhibited.

**Supplemental Figure S9.** Filament convolutedness decreases when formin or myosin activities are inhibited.

**Supplemental Movie S1.** Full time-lapse sequence of actin filament nucleation activities in mock-treated wild-type (*ARP2*) cells.

**Supplemental Movie S2.** Full time-lapse sequence of actin filament nucleation activities in mock-treated *arp2-1* cells.

**Supplemental Movie S3.** Representative actin filament nucleated *de novo* from the cytoplasm (as in Figure 3 A, top row).

**Supplemental Movie S4.** Representative actin filament nucleated from the side of a pre-existing filament (as in Figure 3 A, middle row).

**Supplemental Movie S5.** Representative actin filament nucleated from the end of a pre-existing filament (as in Figure 3 A, bottom row).

**Supplemental Movie S6.** Representative example of filament elongation (as in Figure 4 A).

**Supplemental Movie S7.** Representative example of filament severing (as in Figure 4 A).

**Supplemental Movie S8.** Full time-lapse sequence of actin filament nucleation activities in CK- 666-treated wild-type (*ARP2*) cells.

**Supplemental Movie S9.** Full time-lapse sequence of actin filament nucleation activities in CK- 666-treated *arp2-1* cells.

**Supplemental Movie S10.** Full time-lapse sequence of actin filament nucleation activities in SMIFH2-treated wild-type (*ARP2*) cells.

**Supplemental Movie S11.** Full time-lapse sequence of actin filament nucleation activities in SMIFH2-treated *arp2-1* cells.

**Supplementary Data Set S1.** ANOVA and Chi-square test results and parameters.

## Acknowledgments

We thank Hongbing Luo (Purdue University) for excellent care and maintenance of plant materials, Dan Szymanski (Purdue University) for providing the *arpc2* line, and David Kovar (University of Chicago) and Laurent Blanchoin (Interdisciplinary Research Institute of Grenoble) for helpful comments on the manuscript. We are also grateful to the Arabidopsis Biological Resource Center (Ohio State University) for supplying the *arp2-1, prf1-2, and fh1-2* T-DNA insertion lines. The TIRF microscopy facility was supported, in part, by the Bindley Bioscience Center at Purdue University. This work was funded by a grant from the Physical Biosciences program of the US Department of Energy, Office of Basic Energy Sciences (DE-FG02- 09ER15526) to C.J.S. Conceptualization of experiments, data analysis, and manuscript preparation by C.J.S. were also supported by the EMBRIO Institute, an NSF-Biological Integration Institute under contract no. 2120200.

The authors declare no competing financial interests.

## Author Contributions

L.X. acquired and analyzed data, prepared figures, wrote and revised the manuscript; L.C. acquired and analyzed data; J.L. acquired and analyzed data; C.J.S. conceptualized experiments, acquired funding, analyzed data, wrote and revised the manuscript.

## Notes

### Competing Interest Statement

The authors have declared no competing interest.

### Summary of Updates

New experimental data added with additional mutant lines (e.g. prf1-2, fh1-2), a complementary actin reporter (LifeAct), different tissue type (cotyledon epidermal cells), as well as revised and expanded statistical analyses for all datasets.

