## Supplemental Figures and Tables for "Cooperative actin filament nucleation by the Arp2/3 complex and formins maintains the homeostatic cortical array in Arabidopsis epidermal cells"

### Supplemental Data

**Table S1**. Single actin filament dynamics in wild type (*ARPC2*) and *arpc2* mutant with or without CK-666.

| Stochastic Dynamic Parameters | Genotype & Treatment | | | |
|---|---|---|---|---|
| | *ARPC2* w/ mock | *arpc2* w/ mock | *ARPC2* w/ CK-666 | *arp2* w/ CK-666 |
| Elongation Rate; µm/s | 1.78 ± 0.05 [a] | 1.87 ± 0.07 [b] | 1.87 ± 0.07 [b] | 1.86 ± 0.07 [b] |
| Max. Filament Length; µm | 16.4 ± 0.3 [a] | 17.2 ± 0.4 [b] | 17.2 ± 0.3 [b] | 17.5 ± 0.3 [b] |
| Max. Filament Lifetime; s | 18.2 ± 0.6 [a] | 22.2 ± 0.3 [b] | 21.8 ± 0.3 [b] | 23.2 ± 0.4 [b] |
| Severing Frequency; breaks/mm/s | 8.08 ± 0.18 [a] | 8.03 ± 0.17 [a] | 8.12 ± 0.16 [a] | 7.97 ± 0.16 [a] |

Measurements were taken from epidermal cells in the elongating apical region of 5-d-old dark-grown hypocotyls. Values represent mean ± SE. n ≥ 100 filaments from two biological repeats conducted with independent plant materials (for one biological repeat, 5 filaments were counted in one hypocotyl from at least 10 hypocotyls per treatment/genotype group). Two-way ANOVA with Tukey's post-hoc test, letters [a–b] denote genotypes or treatments that show statistically significant differences with other genotypes or treatments, $P < 0.05$

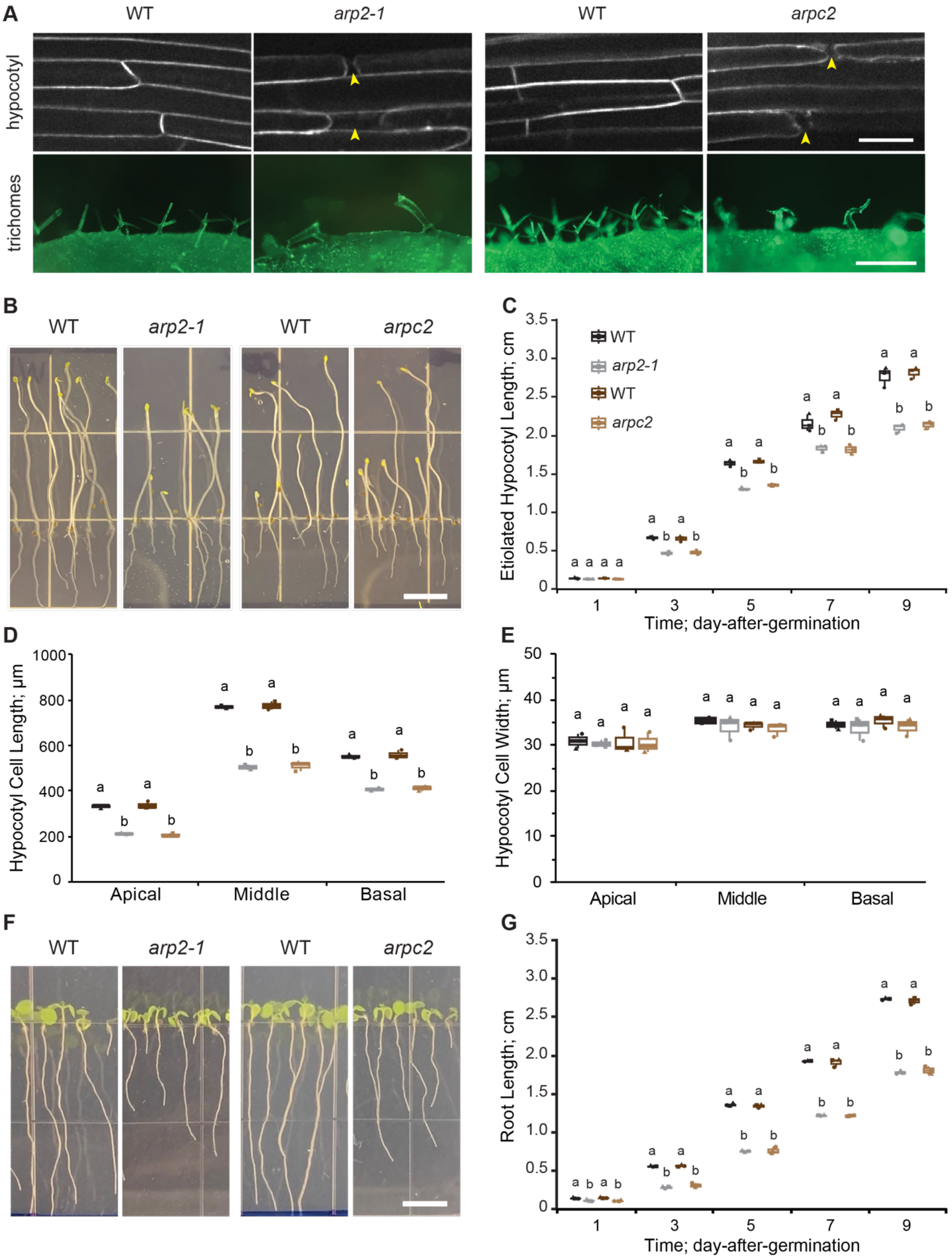
A
WT
arp2-1
WT
arpc2
hypocotyl
trichomes
B
WT
arp2-1
WT
arpc2
C
Etiolated Hypocotyl Length; cm
Time; day-after-germination
WT
arp2-1
WT
arpc2
D
Hypocotyl Cell Length; μm
Apical
Middle
Basal
E
Hypocotyl Cell Width; μm
Apical
Middle
Basal
F
WT
arp2-1
WT
arpc2
G
Root Length; cm
Time; day-after-germination

**Figure S1.** Disruption of the Arp2/3 complex perturbs epidermal cell morphology and inhibits normal plant growth (Supports Figure 1).

**A)** Representative images of epidermal cells from 5-d-old etiolated hypocotyls of homozygous *arp2/3* mutants, *arp2-1* and *arpc2*, and the respective wild-type lines. Cells in *arp2-1* and *arpc2* seedlings had visible cell adhesion defects (yellow arrowheads) at end walls, shown by FM4-64 labeling of the plasma membrane, compared to wild-type sibling lines (top row). Scale bar: 10 µm. Representative images of trichomes from young leaves of 2-week-old plants (bottom row). Trichomes on *arp2-1* and *arpc2* leaves were severely distorted compared to the respective wild-type lines. Scale bar: 8 µm. **B)** Representative images of 5-d-old dark-grown seedlings. Scale bar: 1.0 cm. **C)** Hypocotyls from *arp2-1* and *arpc2* were significantly shorter than the respective wild-type lines over a developmental time series. **D and E**) Quantitative analysis of average length **(D)** and width **(E)** of epidermal cells from the apical, middle, and basal regions of 5-d-old etiolated hypocotyls. **F)** Representative images of 7-d-old light-grown seedlings. Scale bar: 1.0 cm. G, Roots from *arp2-1* and *arpc2* seedlings were significantly shorter than the respective wild-type lines over a developmental time series. For box-and-whisker plots in **(C)** and **(G)**, boxes show the interquartile range (IQR) and the median, and whiskers show the maximum-minimum interval of three biological repeats with independent populations of plants. Individual biological repeats are represented with different shapes ($n = 3$ biological repeats; each data point represents the mean value from at least 50 seedlings per genotype). Letters [a–b] denote groups that show statistically significant differences with other genotypes within the same time point by one-way ANOVA with Tukey's post-hoc test ($P < 0.05$). For box-and-whisker plots in **(D)** and **(E)**, boxes show the interquartile range (IQR) and the median, and whiskers show the maximum-minimum interval of three biological repeats with independent populations of plants. Individual biological repeats are represented with different shapes ($n = 3$ biological repeats; each data point represents the mean value from at least 50 cells from 10 hypocotyls per genotype). Letters [a–b] denote groups that show statistically significant differences with other genotypes within the same region by one-way ANOVA with Tukey's post-hoc test ($P < 0.05$).

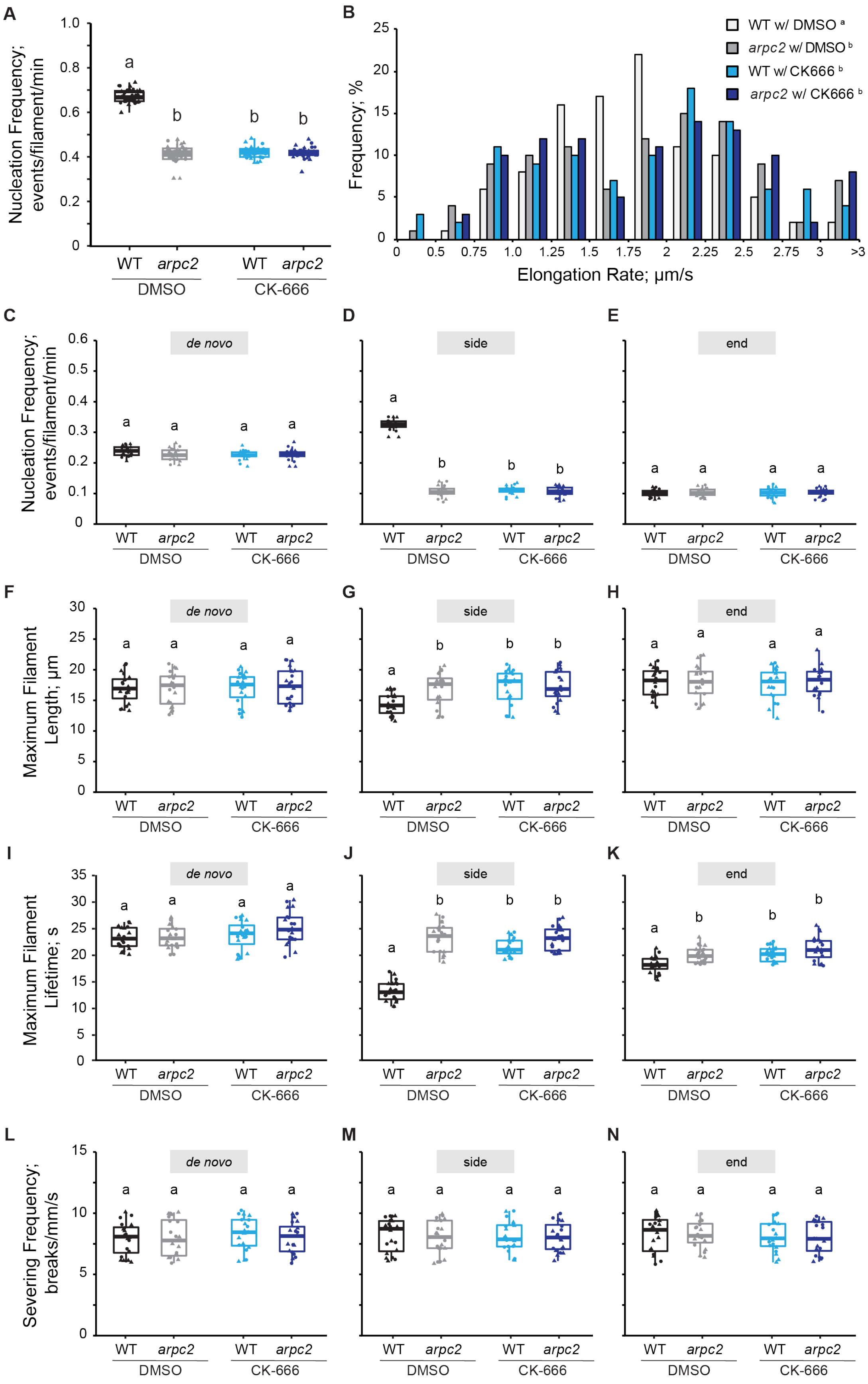
A
Nucleation Frequency; events/filament/min
WT
arpc2
DMSO
CK-666
B
WT w/ DMSO a
arpc2 w/ DMSO b
WT w/ CK666 b
arpc2 w/ CK666 b
Frequency; %
Elongation Rate; µm/s
C
D
E
de novo
side
end
Nucleation Frequency; events/filament/min
F
G
H
Maximum Filament Length; µm
I
J
K
Maximum Filament Lifetime; s
L
M
N
Severing Frequency; breaks/mm/s

**Figure S2.** Actin filament nucleation frequency is decreased in *arpc2* and after treatment of wild type with CK-666 (Support Figure 3).

**A and C–E)** Quantitative analysis of actin filament nucleation frequency. Total nucleation frequency **(A)** in DMSO-treated wild-type cells was significantly higher than DMSO-treated *arpc2*, CK-666-treated wild-type, or CK-666-treated *arpc2* cells. When filament origin events were categorized into *de novo*, side, and end populations **(C–E)**, only the side-branching nucleation events showed a significant reduction in *arpc2* and CK-666-treated cells compared to DMSO-treated wild type. In box-and-whisker plots, boxes show the interquartile range (IQR) and the median, and whiskers show the maximum-minimum interval of two biological repeats with independent populations of plants. Individual biological repeats are represented with different shapes (n = 20 seedlings, 10 seedlings per biological repeat). Letters [a–b] denote groups that show statistically significant differences with other genotypes or treatments by two-way ANOVA with Tukey's post-hoc test ($P < 0.05$). **B)** Quantitative analysis of the population distribution and average elongation rate of actin filaments in hypocotyl epidermal cells. The elongation rate distribution of DMSO-treated wild type had a single peak at 1.25 – 1.75 µm/s, whereas the Arp2/3-inhibited groups had three peaks at 0.75 – 1. 5 µm/s, 2.0 – 2. 5 µm/s, and > 3 µm/s. $n \geq 100$ single filaments from two individual biological repeats (for one biological repeat, 5 single filaments were counted in one hypocotyl from at least 10 hypocotyls per genotype or treatment). Letters [a–b] denote genotypes or treatments that show statistically significant differences with other groups by Chi-squared test, $P < 0.05$. **F–H)** The average maximum length of side-branching filaments in DMSO-treated wild-type cells was significantly shorter than that in Arp2/3-inhibited cells; however, filaments that originated *de novo* or from pre-existing ends did not show any significant difference. **I–K)** The average maximum lifetime of side-branching filaments in DMSO-treated wild-type cells was significantly shorter than that in Arp2/3-inhibited cells, but filaments that originated from pre-existing ends did not show a difference between any genotype or treatment. **L–N)** The severing frequency did not show any significant difference between different genotypes or treatments. For box-and-whisker plots in **(F–N)**, boxes show the interquartile range (IQR) and the median, and whiskers show the maximum-minimum interval of two biological repeats with independent populations of plants. Individual biological repeats are represented with different shapes (n = 20 seedlings, 10 seedlings per biological repeat). Letters [a–b] denote groups that show statistically significant differences with other genotypes or treatments within the same filament nucleation subclass by two-way ANOVA with Tukey's post-hoc test ($P < 0.05$).

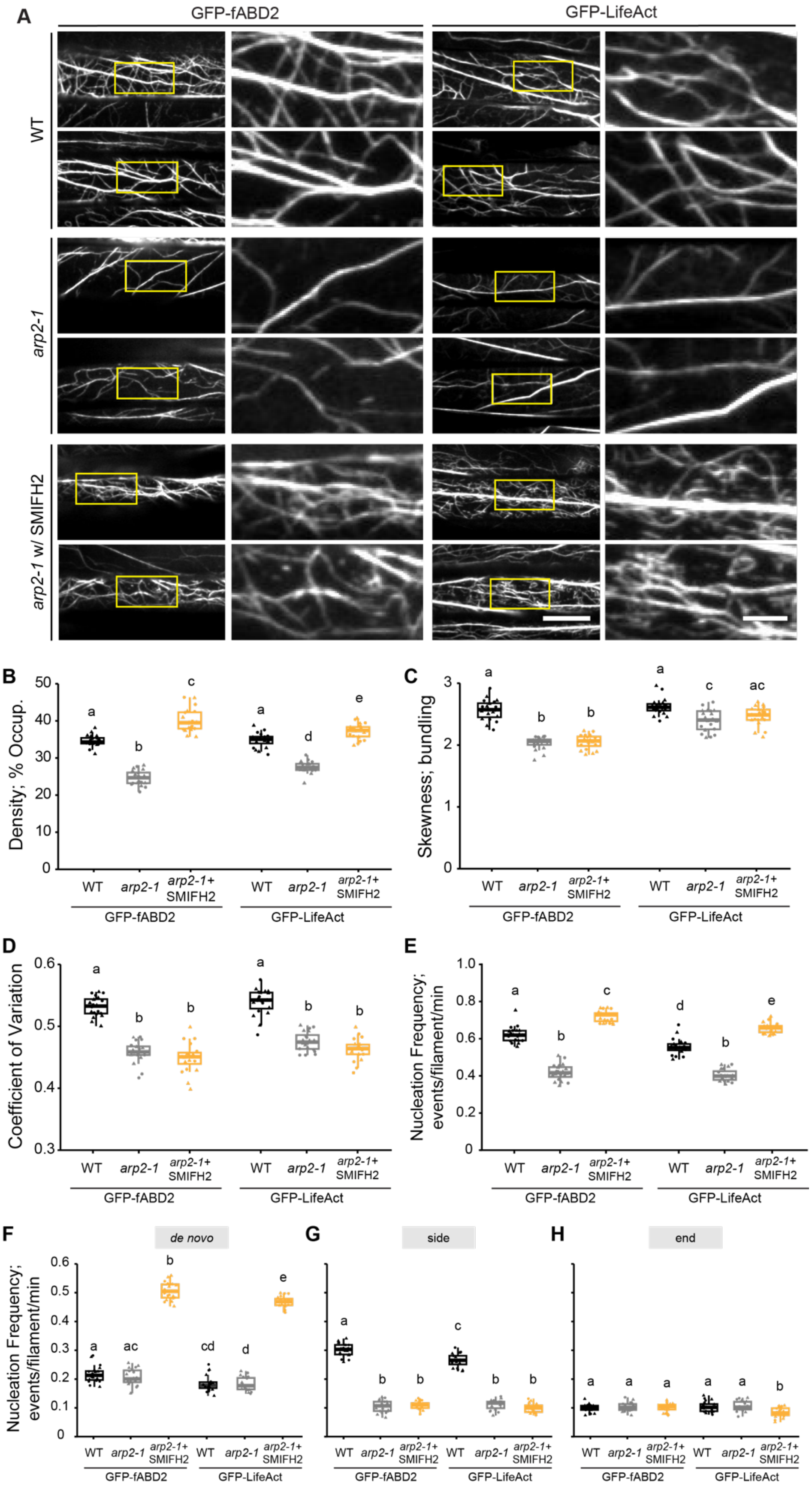
A
GFP-fABD2
GFP-LifeAct
WT
arp2-1
arp2-1 w/ SMIFH2
B
Density; % Occup.
C
Skewness; bundling
D
Coefficient of Variation
E
Nucleation Frequency; events/filament/min
F
de novo
G
side
H
end
WT
arp2-1
arp2-1+ SMIFH2
GFP-fABD2
GFP-LifeAct

**Figure S3.** Two different actin cytoskeleton markers, GFP-fABD2 and GFP-LifeAct, report similar changes in actin array organization and nucleation events regardless of the genotype or treatment (Supports Figures 1 and 2).

**A)** Representative images of epidermal cells from the apical region of 5-d-old etiolated hypocotyls expressing GFP-fABD2 or GFP-LifeAct in wild type or *arp2-1* lines are shown in the left columns. Scale bar: 20 µm. Regions of interest (yellow boxes) were magnified and displayed in the right columns. Scale bar: 5 µm. Hypocotyls were treated with 0.05% DMSO solution or 25 µM SMIFH2 for 5 min prior to imaging. Actin filament arrays in DMSO-treated *arp2-1* cells appeared to be less dense compared to DMSO-treated wild-type cells; however, SMIFH2-treated *arp2-1* cells appeared to have more dense actin arrays compared to DMSO-treated wild-type cells, regardless of the actin cytoskeleton reporter used (GFP-fABD2 or -LifeAct) **B–D)** Quantitative analysis of the percentage of occupancy or density of actin filament arrays **(B)** and the extent of filament bundling as measured by skewness **(C)** and coefficient of variance **(D)** analyses. For cells expressing either GFP-fABD2 or GFP-LifeAct, both the density of actin arrays and the extent of filament bundling in DMSO-treated *arp2-1* cells were significantly decreased but SMIFH2-treated *arp2-1* cells had a significantly increased actin density compared to DMSO-treated wild-type cells. In box-and-whisker plots, boxes show the interquartile range (IQR) and the median, and whiskers show the maximum-minimum interval of two biological repeats with independent populations of plants. Individual biological repeats are represented with different shapes (n = 20 seedlings, 10 seedlings per biological repeat). Letters [a–e] denote groups that show statistically significant differences with other genotypes or treatments by two-way ANOVA with Tukey's post-hoc test ($P < 0.05$). **E–H)** Quantitative analysis of actin filament nucleation frequency, both overall **(E)** and by subclass of origin **(F–H)**. The total nucleation frequency in DMSO-treated *arp2-1* cells expressing either GFP-fABD2 or GFP-LifeAct was significantly reduced compared to the corresponding DMSO-treated wild-type cells. Moreover, similar increased total and *de novo* nucleation frequencies were observed in SMIFH2-treated *arp2-1* cells irrespective of the reporter used. In box-and-whisker plots, boxes show the interquartile range (IQR) and the median, and whiskers show the maximum-minimum interval of two biological repeats with independent populations of plants. Individual biological repeats are represented with different shapes (n = 20 seedlings, 10 seedlings per biological repeat). Letters [a–e] denote groups that show statistically significant differences with other genotypes or treatments by two-way ANOVA with Tukey's post-hoc test ($P < 0.05$).

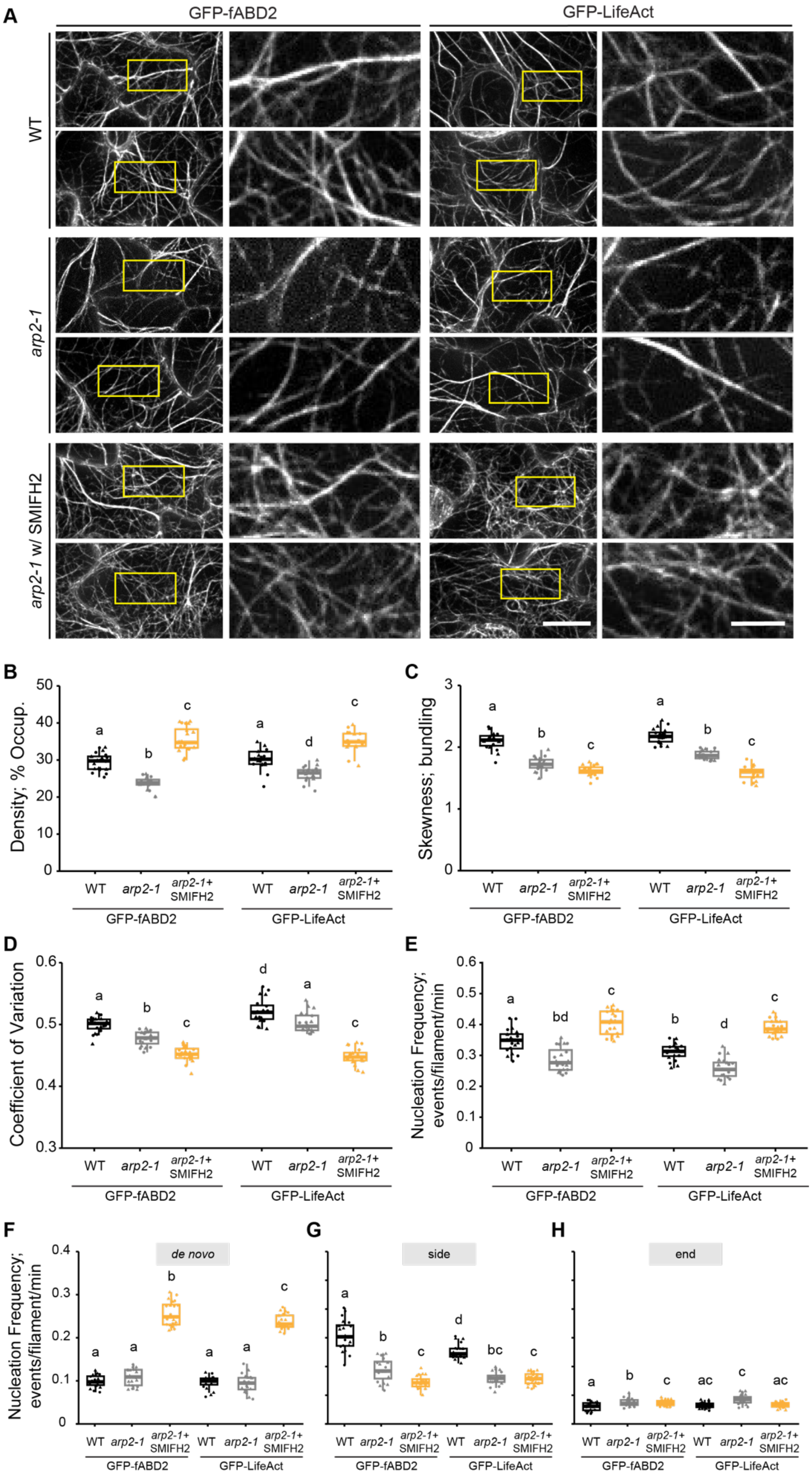

A
GFP-fABD2
GFP-LifeAct
WT
arp2-1
arp2-1 w/ SMIFH2
B
Density; % Occup.
WT arp2-1 arp2-1+ SMIFH2
GFP-fABD2
GFP-LifeAct
C
Skewness; bundling
D
Coefficient of Variation
E
Nucleation Frequency; events/filament/min
F
de novo
G
side
H
end

**Figure S4.** Cotyledon epidermal cells show similar phenotypes of reduced actin filament abundance and decreased nucleation frequency in *arp2-1* mutants expressing either GFP-fABD2 or GFP-LifeAct cytoskeletal reporters (Supports Figures 1 and 2).

**A)** Representative images of adaxial epidermal cells from 5-d-old cotyledons expressing either GFP-fABD2 or GFP-LifeAct in wild type and *arp2-1* lines are shown in the left columns. Scale bar: 10 µm. Regions of interest (yellow box) were magnified and displayed in the right columns. Scale bar: 3 µm. Hypocotyls were treated with 0.05% DMSO solution or 25 µM SMIFH2 for 5 min prior to imaging. Actin filament arrays in DMSO-treated *arp2-1* cells appeared to be less dense compared to DMSO-treated wild-type cells, regardless of which cytoskeletal reporter was expressed in the line. **B–D)** Quantitative analysis of the percentage of occupancy or density of actin filament arrays **(B)** and the extent of filament bundling as measured by skewness **(C)** and coefficient of variance **(D)** analyses. Cells expressing either GFP-fABD2 or GFP-LifeAct showed similar changes in actin array organization between wild type and *arp2-1*. Both the density of actin arrays and the extent of filament bundling in DMSO-treated *arp2-1* cells were significantly decreased but SMIFH2-treated *arp2-1* cells had a significantly increased actin density compared to DMSO-treated wild-type cells, regardless of which cytoskeletal reporter was expressed. In box-and-whisker plots, boxes show the interquartile range (IQR) and the median, and whiskers show the maximum-minimum interval of two biological repeats with independent populations of plants. Individual biological repeats are represented with different shapes (n = 20 seedlings, 10 seedlings per biological repeat). Letters [a–d] denote groups that show statistically significant differences with other genotypes or treatments by two-way ANOVA with Tukey's post-hoc test ($P < 0.05$). **E–H)** Quantitative analysis of actin filament nucleation frequency, both overall **(E)** and by subclass of origin **(F–H)**. Irrespective of the cytoskeletal reporter expressed in the line, the total nucleation frequency in DMSO-treated *arp2-1* cells was significantly reduced compared to DMSO-treated wild-type cells, but the total nucleation frequency of SMIFH2-treated *arp2-1* cells was significantly higher than all other genotypes and treatments and this correlated with increased *de novo* nucleation events. In box-and-whisker plots, boxes show the interquartile range (IQR) and the median, and whiskers show the maximum-minimum interval of two biological repeats with independent populations of plants. Individual biological repeats are represented with different shapes (n = 20 seedlings, 10 seedlings per biological repeat). Letters [a–d] denote groups that show statistically significant differences with other genotypes or treatments (within the same filament nucleation subclass) by two-way ANOVA with Tukey's post-hoc test ($P < 0.05$).

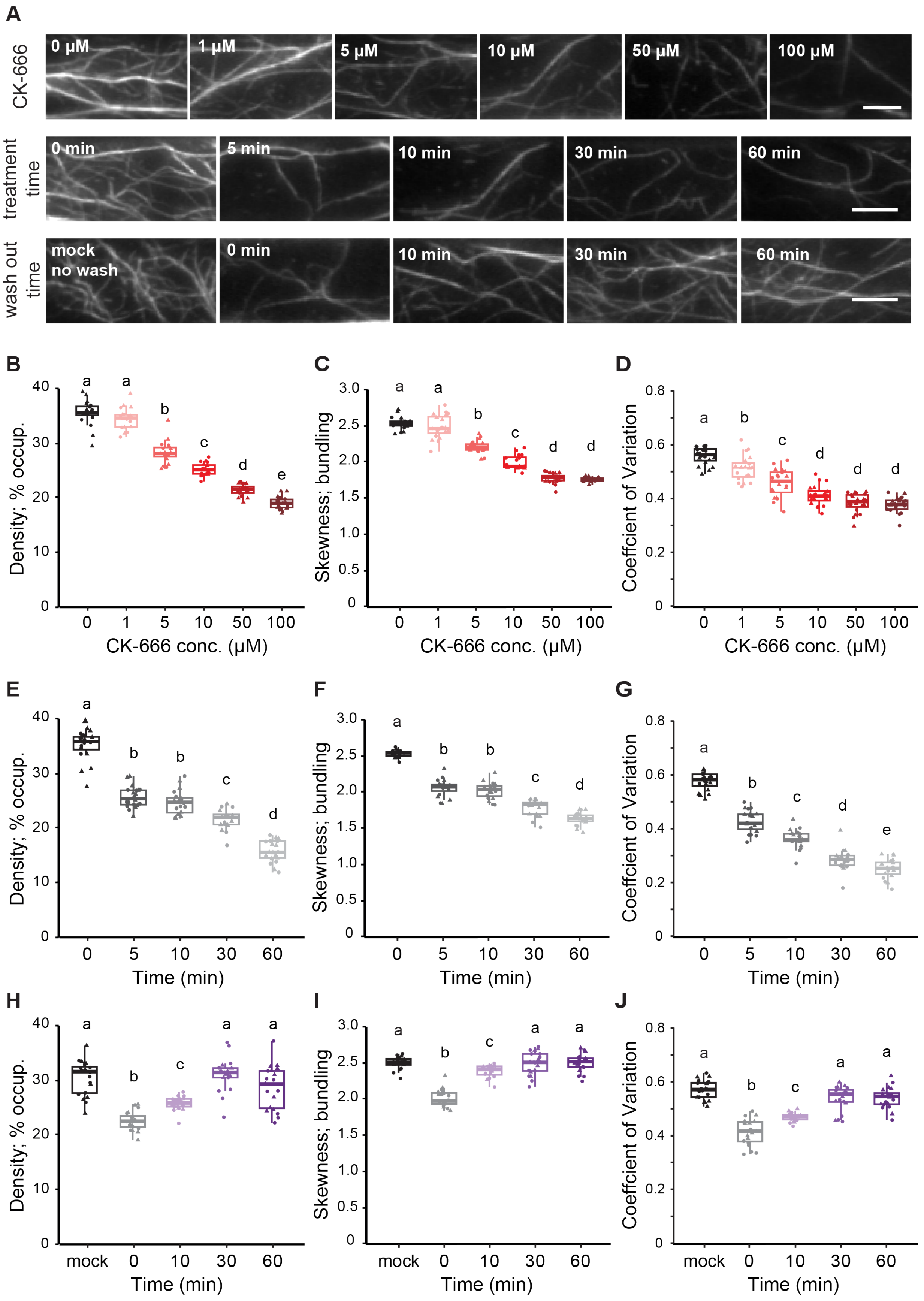
A
CK-666
0 μM
1 μM
5 μM
10 μM
50 μM
100 μM
treatment time
0 min
5 min
10 min
30 min
60 min
wash out time
mock no wash
0 min
10 min
30 min
60 min
B
Density; % occup.
CK-666 conc. (μM)
C
Skewness; bundling
CK-666 conc. (μM)
D
Coeffcient of Variation
CK-666 conc. (μM)
E
Density; % occup.
Time (min)
F
Skewness; bundling
Time (min)
G
Coeffcient of Variation
Time (min)
H
Density; % occup.
Time (min)
I
Skewness; bundling
Time (min)
J
Coeffcient of Variation
Time (min)

**Figure S5.** CK-666 is an acute and reversible chemical inhibitor of the Arp2/3 complex in plant cells (Supports Figures 2 and 4).

**A)** Representative images of actin filaments from epidermal cells of 5-d-old etiolated hypocotyls after treatment with a dose series of CK-666 for 5 min (top), with 10 μM CK-666 for a series of time (middle), or with 10 μM CK-666 for 5 min then washed out with water for a series of time (bottom). Scale bar: 10 μm. **B–J)** Quantitative analysis of the percentage of occupancy or density of actin filament arrays **(B, E, and H)** as well as the extent of filament bundling as measured by skewness **(C, F and I)** and coefficient of variance **(D, G and J)** analyses. Both the density **(B)** and bundling **(C–D)** of actin arrays significantly decreased after treatment with 10 μM CK-666 and treating hypocotyls for 5 min was sufficient to elicit changes in array architecture **(E–G)**. After a 5 min treatment with 10 μM CK-666, the density **(H)** and bundling **(I–J)** of actin arrays returned to normal following wash out for 30 min. In box-and-whisker plots, boxes show the interquartile range (IQR) and the median, and whiskers show the maximum-minimum interval of two biological repeats with independent populations of plants. Individual biological repeats are represented with different shapes (n = 20 seedlings, 10 seedlings per biological repeat). Letters [a–e] denote treatments that show statistically significant differences with other treatments by one-way ANOVA with Tukey's post-hoc test ($P < 0.05$).

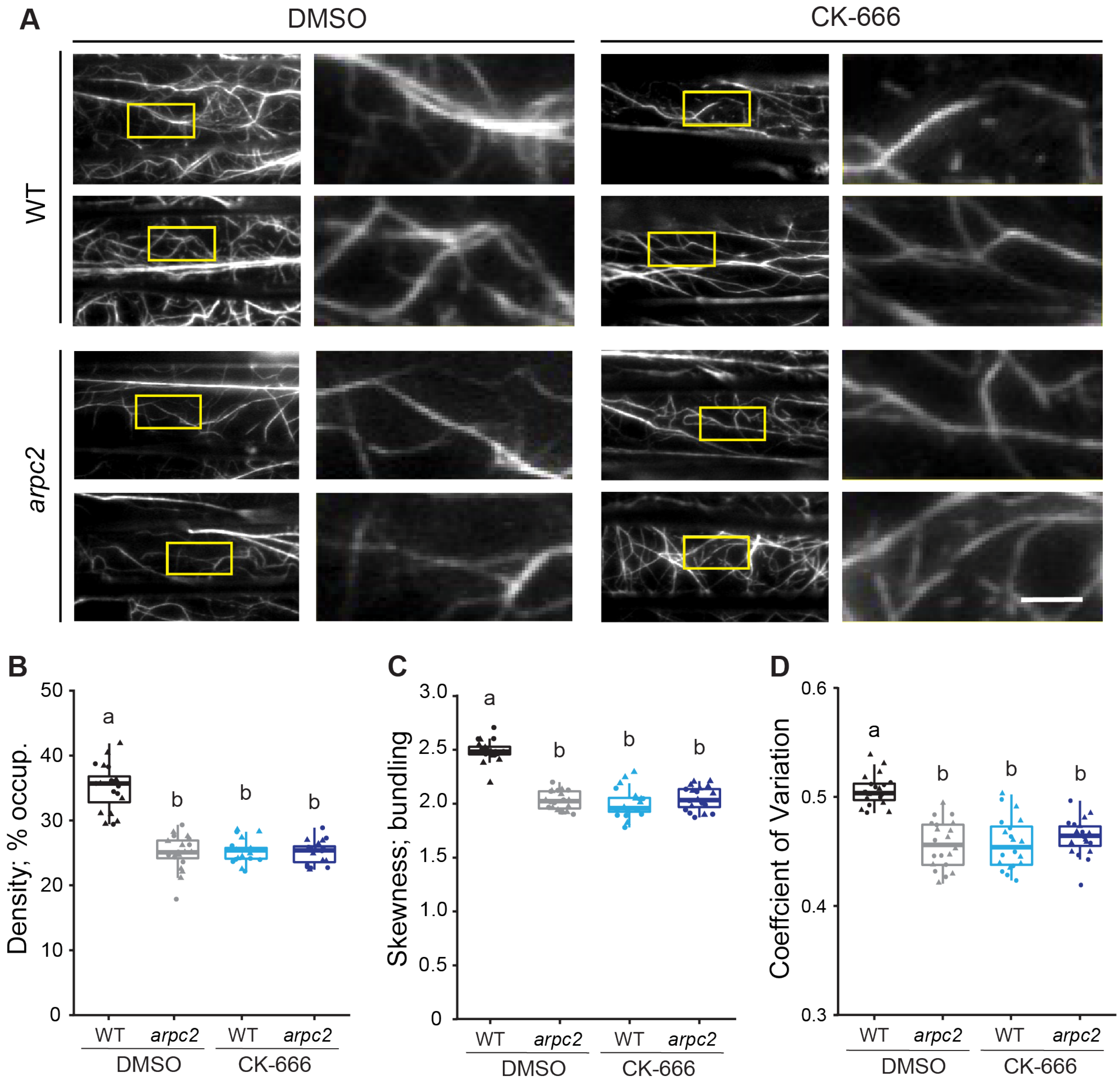


**Figure S6.** Actin array architecture is perturbed in *arpc2* and the reduced filament density and bundling phenotypes are recapitulated by treatment with CK-666 (Supports Figure 4).

**A)** Representative images of epidermal cells from the apical region of 5-d-old etiolated hypocotyls are shown in the left columns. Scale bar: 20 µm. Regions of interest (yellow boxes) were magnified and displayed in the right columns. Scale bar: 5 µm. Hypocotyls were treated with 0.05% DMSO solution or 10 µM CK-666 for 5 min prior to imaging with VAEM. Actin filament arrays in DMSO-treated *arpc2*, CK-666-treated wild-type, and CK-666-treated *arpc2* cells appeared to be less dense and less bundled compared to DMSO-treated wild-type cells. **B–D)** Quantitative analysis of the percentage of occupancy or density of actin filament arrays **(B)** and the extent of filament bundling as measured by skewness **(C)** and coefficient of variance **(D)** analyses. Both the density and bundling of actin arrays in DMSO-treated *arpc2*, CK-666-treated wild type and CK-666-treated *arpc2* cells were significantly decreased compared to DMSO-treated wild-type cells. In box-and-whisker plots, boxes show the interquartile range (IQR) and the median, and whiskers show the maximum-minimum interval of two biological repeats with independent populations of plants. Individual biological repeats are represented with different shapes (n = 20 seedlings, 10 seedlings per biological repeat). Letters [a–b] denote groups that show statistically significant differences with other genotypes or treatments by two-way ANOVA with Tukey's post-hoc test ($P < 0.05$).

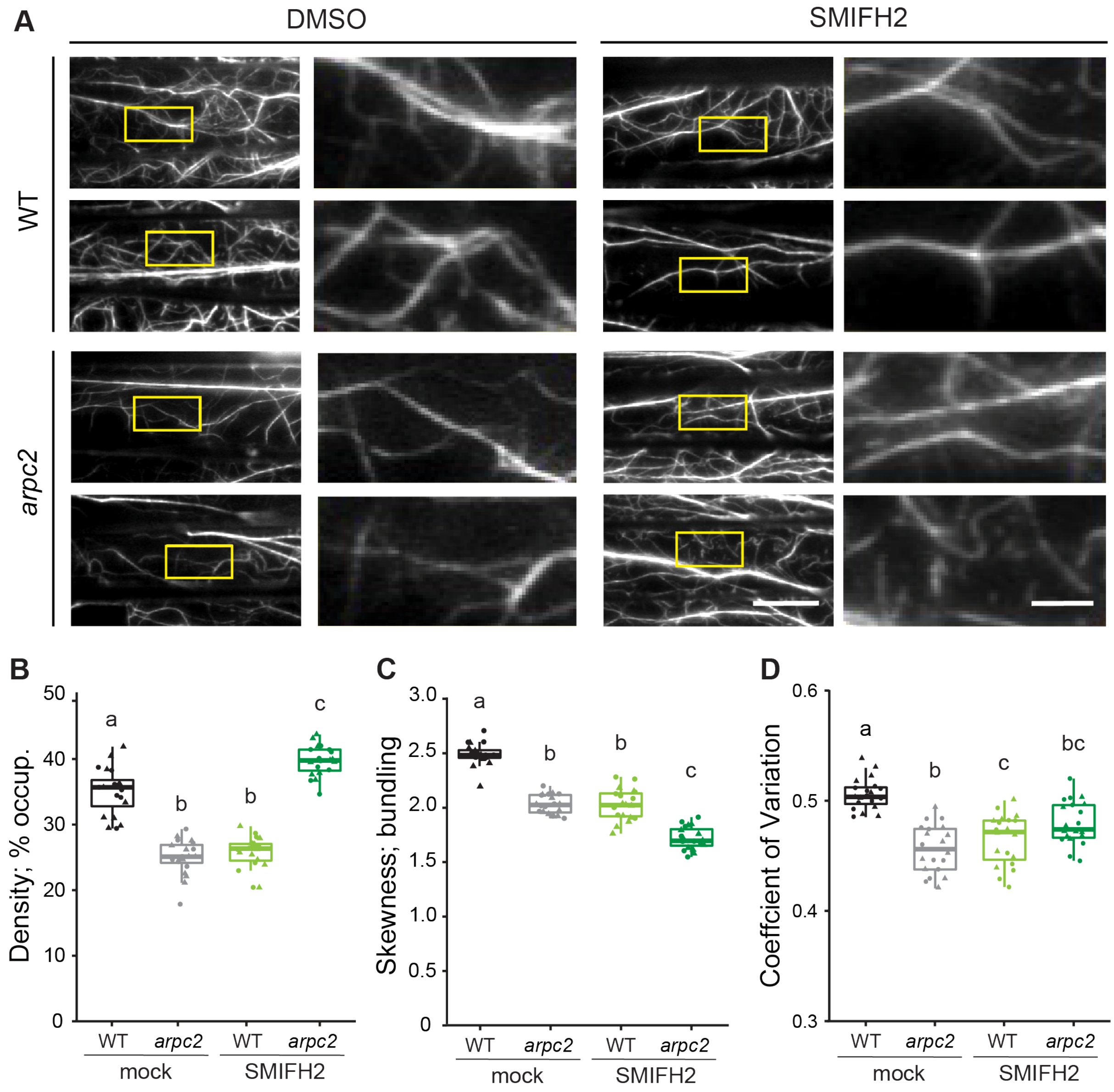


**Figure S7.** Actin filament density increases after treatment of *arpc2* with the formin inhibitor SMIFH2 (Supports Figure 5).

**A)** Representative images of epidermal cells from the apical region of 5-day-old etiolated hypocotyls are shown in the left columns. Scale bar: 20 µm. Regions of interest (yellow boxes) were magnified and displayed in the right columns. Scale bar: 5 µm. Hypocotyls were treated with 0.05% DMSO solution or 25 µM SMIFH2 for 5 min prior to imaging with VAEM. Actin filament arrays in DMSO-treated *arpc2*, SMIFH2-treated wild-type and SMIFH2-treated *arpc2* cells appeared to be less dense and less bundled compared to DMSO-treated wild-type cells. **B–D)** Quantitative analysis of the percentage of occupancy or density of actin filament arrays **(B)** and the extent of filament bundling as measured by skewness **(C)** and coefficient of variance **(D)** analyses. The density of actin arrays in DMSO-treated *arpc2* and SMIFH2-treated wild-type cells was significantly decreased compared to DMSO-treated wild-type; however, SMIFH2-treated *arpc2* cells had significantly increased actin density compared to all other genotypes and treatments **(B)**. Actin arrays in *arpc2* or SMIFH2-treated cells were significantly less bundled compared to DMSO-treated wild-type cells **(C–D)**. In box-and-whisker plots, boxes show the interquartile range (IQR) and the median, and whiskers show the maximum-minimum interval of two biological repeats with independent populations of plants. Individual biological repeats are represented with different shapes (n = 20 seedlings, 10 seedlings per biological repeat). Letters [a–c] denote groups that show statistically significant differences with other genotypes or treatments by two-way ANOVA with Tukey's post-hoc test ($P < 0.05$).

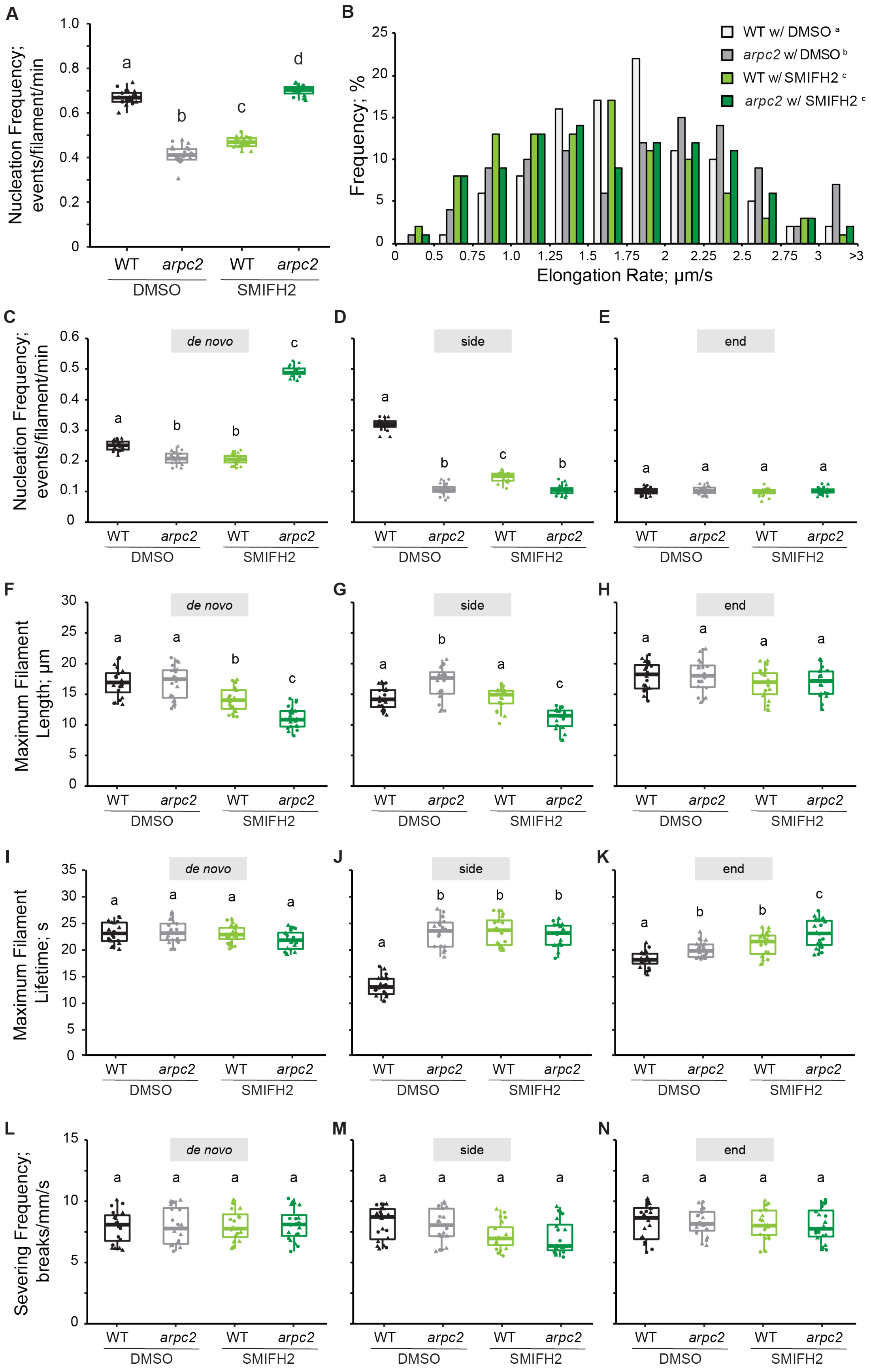

A
Nucleation Frequency; events/filament/min
1.0
0.8
0.6
0.4
0.2
0
a
b
c
d
WT
arpc2
WT
arpc2
DMSO
SMIFH2
B
WT w/ DMSO a
arpc2 w/ DMSO b
WT w/ SMIFH2 c
arpc2 w/ SMIFH2 c
Frequency; %
25
20
15
10
5
0
0
0.5
0.75
1.0
1.25
1.5
1.75
2
2.25
2.5
2.75
3
>3
Elongation Rate; µm/s
C
D
E
Nucleation Frequency; events/filament/min
0.6
0.5
0.4
0.3
0.2
0.1
0
de novo
side
end
F
G
H
Maximum Filament Length; µm
30
25
20
15
10
5
0
I
J
K
Maximum Filament Lifetime; s
35
30
25
20
15
10
5
0
L
M
N
Severing Frequency; breaks/mm/s
15
10
5
0
WT
arpc2
WT
arpc2
DMSO
SMIFH2

**Figure S8.** Overall and *de novo* filament nucleation increase when ARPC2 and formin activity are simultaneously inhibited (Supports Figure 6).

**A and C–E)** Quantitative analysis of actin filament nucleation frequency, both overall **(A)** and by the subclass of origin **(C–E)**. The total nucleation frequency in either DMSO-treated *arpc2* or SMIFH2-treated wild-type cells was significantly reduced compared to DMSO-treated wild-type cells. However, the total nucleation frequency of SMIFH2-treated *arpc2* cells was significantly higher than all other genotypes and treatments **(A)**, and this correlated with increased *de novo* nucleation events **(C)**. In box-and-whisker plots, boxes show the interquartile range (IQR) and the median, and whiskers show the maximum-minimum interval of two biological repeats with independent populations of plants. Individual biological repeats are represented with different shapes (n = 20 seedlings, 10 seedlings per biological repeat). Letters [a–c] denote groups that show statistically significant differences with other genotypes or treatments (within the same filament nucleation subclass) by two-way ANOVA with Tukey's post-hoc test ($P < 0.05$). **B)** Analysis of the population distribution of actin filament elongation rates. The elongation rate distribution of filaments in DMSO-treated wild-type cells had a single peak at 1.25 – 1.75 µm/s, the DMSO-treated *arpc2* had three peaks at 0.75 – 1.25 µm/s, 1.75 – 2.5 µm/s and > 3 µm/s, the SMIFH2-treated wild-type had a peak at 0.75 – 1. 75 µm/s, but SMIFH2-treated *arpc2* had peaks at 1.0 – 1.5 µm/s and 2 – 2.5 µm/s. n ≥ 100 single filaments from two individual biological repeats (for one biological repeat, 5 single filaments were counted in one hypocotyl from at least 10 hypocotyls per genotype or treatment). Letters [a–c] denote genotypes or treatments that show statistically significant differences with other groups by Chi-squared test, $P < 0.05$. **F–H)** The average maximum length of filaments that originated *de novo* or from side-branching events in SMIFH2-treated *arpc2* was significantly shorter than that in other cells, but filaments that originated from pre-existing ends did not show a difference between any genotype or treatment. **I–K)** The average maximum lifetime of side-branching filaments in DMSO-treated *arpc2*, SMIFH2-treated *arpc2* and SMIFH2-treated wild-type cells was significantly longer than the ones from DMSO-treated wild-type cells; however, filaments that originated *de novo* did not show any significant difference, and filaments that elongated from pre-existing ends in DMSO-treated wild-type cells were slightly shorter than SMIFH2-treated cells. **L–N)** The severing frequency did not show any significant difference between different genotypes or treatments. For box-and-whisker plots in **(F–N)**, boxes show the interquartile range (IQR) and the median, and whiskers show the maximum-minimum interval of two biological repeats with independent populations of plants. Individual biological repeats are represented with different shapes (n = 20 seedlings, 10 seedlings per biological repeat). Letters [a–c] denote groups that show statistically significant differences with other genotypes or treatments (within the same filament nucleation subclass) by two-way ANOVA with Tukey's post-hoc test ($P < 0.05$).

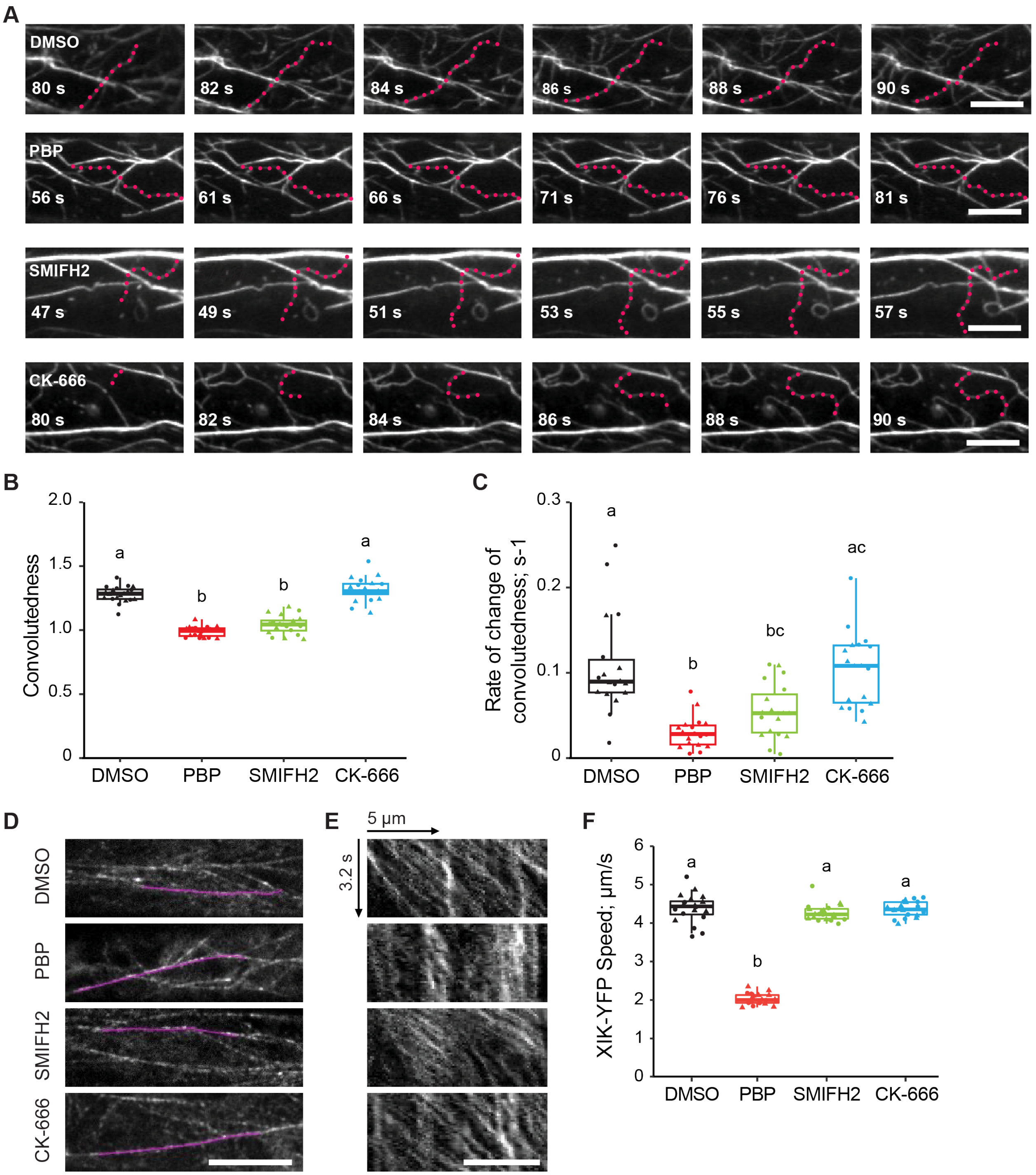


**Figure S9.** Filament convolutedness decreases when formin or myosin activities are inhibited (Supports Figures 5–7).

**A)** Representative time-lapse series of actin filaments undergoing continuous growth, undulations, and buckling. The highlighted filaments (magenta dots) are from Col-0 hypocotyl epidermal cells expressing GFP-fABD2 after treatment with 0.1% DMSO solution, 10 µM PBP, 25 µM SMIFH2, or 10 µM CK-666 for 15 min prior to imaging with VAEM. Scale bar: 10 µm. B–C) Quantitative analyses of filament convolutedness (B) and the rate of change of convolutedness (C) of actin filaments. Both PBP and

SMIFH2 treatments significantly reduced the filament convolutedness and the rate of change of convolutedness compared to the DMSO treatment. However, CK-666 treatment did not have any significant influence on the filament convolutedness. **D)** Representative single-timepoint images of the cortical cytoplasm in epidermal cells expressing myosin XIK-YFP from hypocotyls after treatment with 0.1% DMSO, 10 μM PBP, 25 μM SMIFH2, and 10 μM CK-666 for 15 min prior to imaging with spinning-disk confocal microscopy (SDCM) using an 80-ms interval for 41 frames. Scale bar: 10 μm. **E)** Kymographs of the purple tracks in **(D)** show the movement of the myosin XIK-YFP particles over a 3.2 s time span. Scale bar: 5 μm. **F)** Quantitative analysis of myosin XIK-YFP speed. The motility of YFP-XIK was not influenced in SMIFH2- or CK-666-treated cells compared to DMSO-treated cells, but the PBP-treated cells had significantly reduced YFP-XIK speed. In the box-and-whisker plot, boxes show the interquartile range (IQR) and the median, and whiskers show the maximum-minimum interval of two biological repeats with independent populations of plants. Individual biological repeats are represented with different shapes (n = 20 seedlings, 10 seedlings per biological repeat). Letters [a–b] denote treatments that show statistically significant differences with other treatments by one-way ANOVA with Tukey's post-hoc test ($P < 0.05$).

### Supplemental Movies

**Movie S1**. Full time-lapse sequence of actin filament nucleation activities in DMSO-treated wild-type (*ARP2*) cells. Scale bar: 10 µm. Square, *de novo* filament nucleation; circle, side filament nucleation; triangle, end filament nucleation. Time-lapse images were collected at 1-s intervals and played back at 4 fps. The total elapsed time is 100 s.

**Movie S2**. Full time-lapse sequence of actin filament nucleation activities in DMSO-treated *arp2-1* cells. Scale bar: 10 µm. Square, *de novo* filament nucleation; circle, side filament nucleation; triangle, end filament nucleation. Time-lapse images were collected at 1-s intervals and played back at 4 fps. The total elapsed time is 100 s.

**Movie S3**. Representative actin filament nucleated *de novo* from the cytoplasm (as in Fig. 2 A, top row). Scale bar: 5 µm. Magenta dots, new growing filament; green arrowheads, nucleation site. Time-lapse images were collected at 1-s intervals and played back at 3 fps. The total elapsed time is 20 s.

**Movie S4**. Representative actin filament nucleated from the side of a pre-existing filament (as in Fig. 2 A, middle row). Scale bar: 5 µm. Blue dots, pre-existing filament; magenta dots, new growing filament; green arrowheads, nucleation site. Time-lapse images were collected at 1-s intervals and played back at 3 fps. The total elapsed time is 20 s.

**Movie S5**. Representative actin filament nucleated from the end of a pre-existing filament (as in Fig. 2 A, bottom row). Scale bar: 5 µm. Blue dots, pre-existing filament; magenta dots, new growing filament; green arrowheads, nucleation site. Time-lapse images were collected at 1-s intervals and played back at 3 fps. The total elapsed time is 20 s.

**Movie S6**. Representative example of filament elongation (as in Fig. 3 A). Scale bar: 10 µm. Magenta dots, actin filament; green arrowhead, nucleation site; white asterisk, elongating filament end. Time-lapse images were collected at 1-s intervals and played back at 1 fps. The total elapsed time is 7 s.

**Movie S7**. Representative example of filament severing (as in Fig. 3 A). Scale bar: 10 µm. Magenta dots, actin filament; yellow arrows, severing events. Time-lapse images were collected at 1-s intervals and played back at 1 fps. The total elapsed time is 6 s.

**Movie S8**. Full time-lapse sequence of actin filament nucleation activities in CK-666-treated wild-type (*ARP2*) cells. Scale bar: 10 µm. Square; *de novo* filament nucleation; circle, side filament nucleation; triangle, end filament nucleation. Time-lapse images were collected at 1-s intervals and played back at 4 fps. The total elapsed time is 100 s.

**Movie S9**. Full time-lapse sequence of actin filament nucleation activities in CK-666-treated *arp2-1* cells. Scale bar: 10 µm. Square, *de novo* filament nucleation; circle, side filament nucleation; triangle, end filament nucleation. Time-lapse images were collected at 1-s intervals and played back at 4 fps. The total elapsed time is 100 s.

**Movie S10**. Full time-lapse sequence of actin filament nucleation activities in SMIFH2-treated wild-type (*ARP2*) cells. Scale bar: 10 µm. Square, *de novo* filament nucleation; circle, side filament nucleation; triangle, end filament nucleation. Time-lapse images were collected at 1-s intervals and played back at 4 fps. The total elapsed time is 100 s.

**Movie S11**. Full time-lapse sequence of actin filament nucleation activities in SMIFH2-treated *arp2-1* cells. Scale bar: 10 µm. Square, *de novo* filament nucleation; circlet side filament nucleation; triangle, end filament nucleation. Time-lapse images were collected at 1-s intervals and played back at 4 fps. The total elapsed time is 100 s.
